# Inferring cell differentiation maps from lineage tracing data

**DOI:** 10.1101/2024.09.09.611835

**Authors:** Palash Sashittal, Richard Y. Zhang, Benjamin K. Law, Alexander Strzalkowski, Henri Schmidt, Adriano Bolondi, Michelle M. Chan, Benjamin J. Raphael

**Affiliations:** Dept. of Computer Science, Princeton University, Princeton; 08544 NJ, USA; Lewis-Sigler Institute for Integrative Genomics, Princeton University, Princeton; 08544 NJ, USA; Dept. of Molecular Biology, Princeton University, Princeton; 08544 NJ, USA; Dept. of Genome Regulation, Max Planck Institute for Molecular Genetics; 14195 Berlin, Germany

## Abstract

During development, mulitpotent cells differentiate through a hierarchy of increasingly restricted progenitor cell types until they realize specialized cell types. A cell differentiation map describes this hierarchy, and inferring these maps is an active area of research spanning traditional single marker lineage studies to data-driven trajectory inference methods on single-cell RNA-seq data. Recent high-throughput lineage tracing technologies profile lineages and cell types at scale, but current methods to infer cell differentiation maps from these data rely on simple models with restrictive assumptions about the developmental process. We introduce a mathematical framework for cell differentiation maps based on the concept of potency, and develop an algorithm, *Carta*, that infers an optimal cell differentiation map from single-cell lineage tracing data. The key insight in *Carta* is to balance the trade-off between the complexity of the cell differentiation map and the number of unobserved cell type transitions on the lineage tree. We show that *Carta* more accurately infers cell differentiation maps on both simulated and real data compared to existing methods. In models of mammalian trunk development and mouse hematopoiesis, *Carta* identifies important features of development that are not revealed by other methods including convergent differentiation of specialized cell types, progenitor differentiation dynamics, and the refinement of routes of differentiation via new intermediate progenitors.

**Code availability:** *Carta* software is available at https://github.com/raphael-group/CARTA

## 1 Main

Organismal development occurs via the differentiation of cells through a hierarchy of *progenitor cell types*, each with progressively restricted potential, ultimately leading to specialized cell types. The *cell differentiation map* describes this hierarchy, including all progenitor and specialized cell types and the transitions between these cell types. Deriving cell differentiation maps – of tissues, organs, or complete organisms – is a key challenge in developmental biology.

The traditional method to derive cell differentiation maps involves manual lineage tracing that directly tracks cell division and differentiation during development [1–5]. A notable milestone using this approach was the derivation of the complete differentiation map of the 671 cells of *Caenorhabditis elegans* using time-lapse microscopy [6]. However, such a direct observational approach is not feasible for more complex organisms, such as mice or humans, that contain trillions of cells and develop in utero.

More recently, single-cell RNA sequencing, which measures the transcriptomes of individual cells, has allowed investigation of cell differentiation maps at scale [7–12]. Cell differentiation maps are derived from this data using trajectory inference methods that attempt to infer branching structures and pseudotimes underlying dynamic differentiation processes from transcriptomes measured at one or a small number of timepoints [13–23]. These methods rely on several limiting assumptions that hinder their ability to reconstruct precise cellular relationships, particularly the assumption that all progenitor cell types along the differentiation hierarchy are observed in the data [24, 25].

Recent advances in genome editing and single-cell sequencing have enabled high-throughput lineage tracing of cells in complex developmental systems [26–29]. In these technologies, heritable barcodes are induced in dividing cells using genome editing tools such as CRISPR-Cas9, providing markers of cell divisions. The barcodes can either be introduced at specific stages of development [30–33] or dynamically through a continuous process as cells divide and differentiate [34–39]. Single-cell RNA sequencing simultaneously measures barcodes (revealing the lineage of cells) and gene expression (revealing cell types) for thousands of individual cells as the system develops [40–46]. These barcoding systems offer the scalability to investigate development in complex organisms but have limited resolution compared to exhaustive microscopy methods such as those used for *C. elegans*. Thus, with these technologies one does not typically observe the differentiation decisions of each dividing cell during development.

Current approaches to infer cell differentiation maps from single-cell RNA sequencing or lineage tracing data are based on two opposing assumptions about the number of progenitor cell types that exist in the developmental system. First, trajectory inference-based methods assume that *all* progenitor cell types are observed in the data [24, 25]. On the opposite extreme, other recent studies [28, 39, 47] use distance-based heuristics calculated from lineages that implicitly assume that the cell differentiation map is a binary tree and consequently the number of progenitor cell types is exactly one less than the number of observed cell types. Neither of these assumptions is likely to be true in practice; for example, early transient progenitor cell types that arise long before cell collection are likely unobserved, and the cell differentiation map is not always a tree due to phenomena such as alternate routes of differentiation to cell types (convergent differentiation) [28].

We introduce a mathematical model of cell differentiation maps and derive an algorithm *Carta* that infers an optimal cell differentiation map from single-cell lineage tracing data. We represent a cell differentiation map by a directed acyclic graph whose vertices are cell types and whose edges represent transitions (differentiation events) between cell types that occur during development. Importantly, *Carta* does not assume that all progenitor cell types are measured at the time of the experiment. Instead, we introduce the concept of a *potency* set, defining unobserved progenitors by the cell types of their descendants. Using the concept of potency, we demonstrate that there are two competing objectives when inferring a cell differentiation map from lineage tracing data: the complexity of the cell differentiation map and the *discrepancy* between transitions in the map and the cell lineage tree. *Carta* quantifies the trade-off between these objectives and computes an optimal differentiation map for any number of progenitor cell types, providing a rigorous framework to evaluate different hypotheses about cell differentiation maps.

We demonstrate the ability of *Carta* to infer interpretable cell differentiation maps that recapitulate established developmental trajectories. On simulated cell differentiation maps and lineage tracing data, *Carta* more accurately reconstructs the underlying cell differentiation maps compared to existing methods. In an *in vitro* model for mammalian trunk development [29, 48, 49], *Carta* infers a cell differentiation map that provides insights into the differentiation dynamics of neuro-mesodermal progenitors (NMPs) into somitic and neural tube lineages that are not revealed under the restricted frameworks of existing methods. On lineage tracing data from a mouse hematopoiesis model [30], *Carta* infers a cell differentiation map that better recapitulates the established differentiation of hematopoiesis and also has stronger agreement with gene expression compared to existing methods. *Carta* provides a rigorous quantitative framework to derive cell differentiation maps that extends beyond the restrictions of existing methods and provides opportunities to better understand development in a variety of contexts.

## 2 Results

### 2.1 *Carta*: a computational model for Cell Differentiation Mapping

*Carta* infers an optimal cell differentiation map from cell lineage tree(s) by accounting for ambiguities in these trees resulting from sampling and other inherent limitations in data from current lineage tracing technologies (Figure 1a,b). The inputs to *Carta* are *m cell lineage trees* T := {*T*_1_, …, *T*_*m*_}, with each tree *T*_*i*_ describing the cell division history of a distinct biological replicate of the same developmental system. The leaves of each tree correspond to the sequenced cells, the internal vertices represent the ancestral cells, and the edges indicate cell divisions (Figure 1a). Each leaf is labeled by a cell type – typically derived from single-cell RNA sequencing data – but the internal vertices are unlabeled since the cell types of these cells are not measured. Let *S* be the set of *observed cell types*, i.e. the set of cell types that label the leaves of 𝒯.

**Figure 1:**
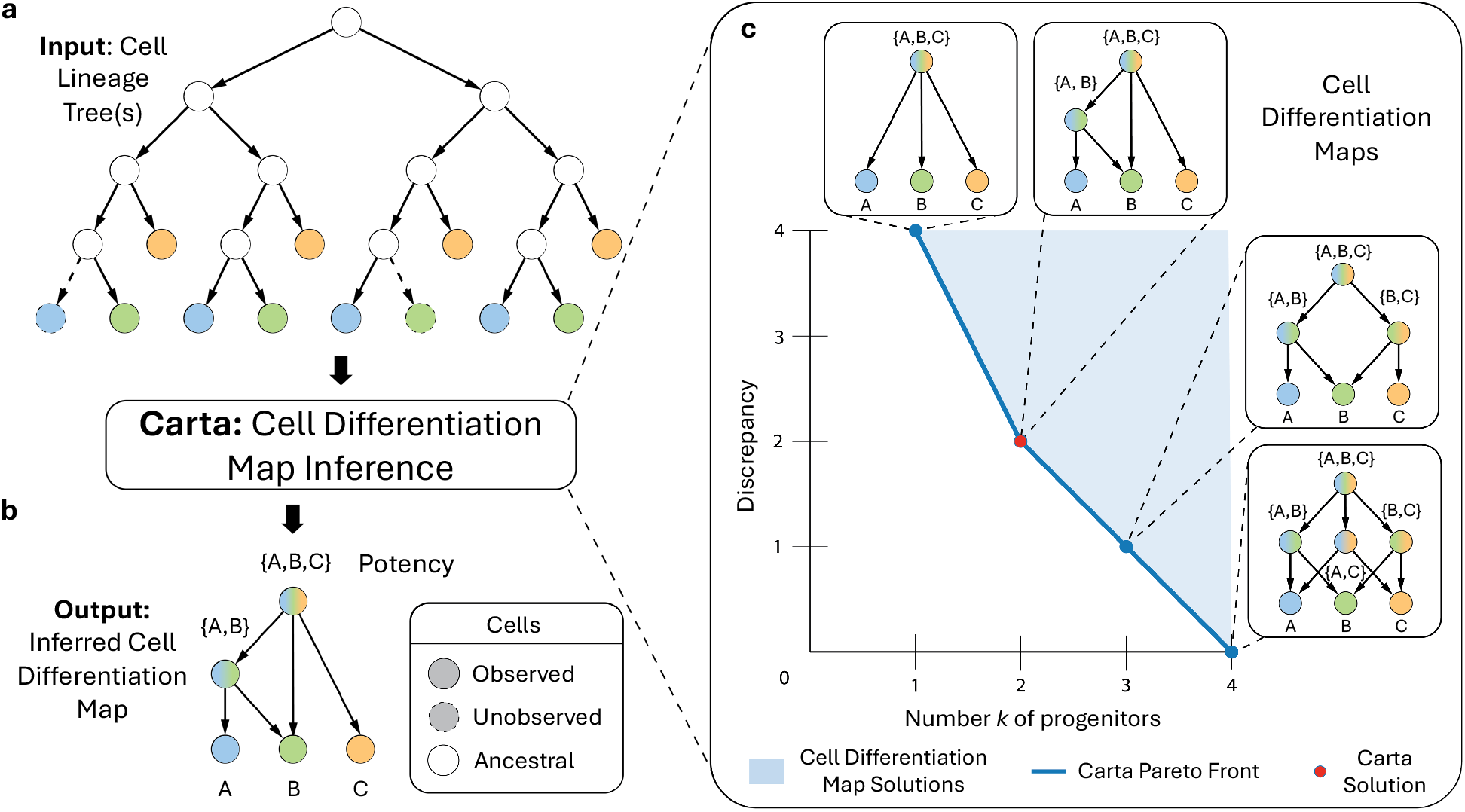
Cell differentiation mapping from lineage tracing data using *Carta*. (a) The input to *Carta* is one or more cell lineage trees, whose leaves are labeled by the measured cell type (labeled *A, B, C*) of the sequenced cells. Typically, some cells that are present at the time of the experiment are not sampled (denoted by dotted lines). (b) *Carta* infers a cell differentiation map that describes the progenitor cell types – represented as a *potency* set – and cell type transitions that occurred during development. (c) *Carta* quantifies the trade-off between the number *k* of progenitor cell types in the cell differentiation map and its *discrepancy* with the cell lineage trees by computing the *Pareto front* of optimal solutions. A cell differentiation map with the optimal number *k*^*^ of progenitors is chosen by identifying an *elbow* of the Pareto front.

A cell differentiation map *F* is a directed graph where the vertices represent cell types and the edges describe the cell type transitions that occurred during development. Given a cell lineage tree with all cells labeled by their cell type, the cell differentiation map is determined by the cell type transitions that occur along the edges of the tree. However, since the cell type of the ancestral cells in the cell lineage trees 𝒯 are not observed, the trees do not directly reveal the cell differentiation map. Thus, inferring a cell differentiation map from lineage tree(s) requires examination of different labelings of the ancestral cells by cell types.

An additional complication in the inference of the cell differentiation map is that the ancestral cells may have cell types that are *not* observed in the lineage tracing data; i.e. the cell type of an ancestral cell may not be observed at the leaves of any tree *T*_*i*_. A key observation in *Carta* is that these unobserved progenitor cell types are described by the set of observed cell types that the progenitor can differentiate into. Namely, each unobserved progenitor cell type corresponds to a *potency* set which contains the observed cell types that are possible future descendants of this cell type. Formally, if *S* is the set of observed cell types, then the potency of a progenitor cell type is a subset of *S*. For instance, the totipotent cell – a progenitor that can differentiate into any observed cell type – has potency *S*, while an observed cell type *t* has potency {*t*}. We define a cell differentiation map *F*_*S*_ for the set *S* of observed cell types to be a directed graph whose vertices represent observed cell types and unobserved progenitors – and are labeled by either an elements of *S or* a subset of *S* – and whose edges represent cell type transitions that occurred during development (Figure 1b).

Multiple cell differentiation maps with varying number *k* of progenitor cell types can explain the development of a set *S* of observed cell types. For example, suppose there are three observed cell types *S* = {A, B, C} (Figure 1a). The simplest cell differentiation map contains *k* = 1 progenitor cell types, namely the totipotent cell with potency *S*, with outgoing edges to vertices representing each of the observed cell types (Figure 1c), top. On the other extreme, the most complex cell differentiation map may contain all possible subsets of *S*, leading to a map with 2^|*S*|^ − |*S*| − 1 progenitor cell types, or *k* = 4 in this example (Figure 1c, bottom). In order to evaluate these differing hypotheses about cellular differentiation, one needs a metric to assess the *fit* between a cell differentiation map and the cell lineage trees.

We introduce the *discrepancy score D*(𝒯, *F*_*S*_), a metric that evaluates the *fit* between a candidate cell differentiation map *F*_*S*_ and a collection 𝒯 of cell lineage trees. *D*(𝒯, *F*_*S*_) relies on a labeling ℓ of the ancestral cells (internal vertices of the cell lineage trees) by potencies of cell types (vertices of the cell differentiation map *F*_*S*_). A *discrepancy* occurs when a cell type in the labeled potency of an ancestral cell is not observed in its leaf descendants. The *discrepancy score D*(𝒯, *F*_*S*_) is the minimum number of such discrepancies when the ancestral cells in 𝒯 are optimally labeled by potencies in *F*_*S*_ (see Methods Section 4.1 for details). A lower discrepancy score indicates a better fit between the cell differentiation map and the cell lineage trees 𝒯 under the assumption that the cells in 𝒯 follow the routes of differentiation in the map.

Solely choosing the cell differentiation map with minimal discrepancy may not lead to accurate inference of the true map due to sampling limitations of current lineage tracing technologies. Specifically, lineage tracing technologies have limited throughput and thus all the cell types in the potency of each ancestral cell may not be observed in the cell lineage trees. For instance, an ancestral cell whose true potency is {A, B, C} may have no descendants with cell type A due to those cells not being sampled (Figure 1a). This leads to an ancestral cell with an observed potency of {B, C}, which does not match any of the progenitor cell types in the true cell differentiation map. As such, minimizing the discrepancy over all possible differentiation maps may lead to the inference of complex cell differentiation maps with large number of progenitors and cell type transitions, several of which may be false positives.

Our algorithm, *Carta*, infers a cell differentiation map *F*_*S*_ from cell lineage trees T by balancing the trade-off between the complexity of *F*_*S*_ and its discrepancy score *D*(T, *F*_*S*_). We characterize the complexity of cell differentiation maps by the number *k* of progenitor cell types. The least complex cell differentiation map (*k* = 1) has a single totipotent progenitor cell type that can differentiate into any observed cell type. However, this map will typically have a very high discrepancy score (Figure 1c, upper left). On the other extreme, one can always find a cell differentiation map with minimum discrepancy *D*(𝒯, *F*_*S*_) = 0, but this map will often have a large number *k* of progenitor cell types, several of which may be false positives (Figure 1c, bottom right). *Carta* solves the Cell Differentiation Map Inference Problem (Problem 4.2 in Methods Section 4.1), deriving a cell differentiation map *F*_*S*_ with minimum discrepancy *D*(𝒯, *F*_*S*_) for each number *k* of progenitor cell types. These solutions give the *Pareto front* of optimal solutions; i.e. there are no cell differentiation maps that have *both* fewer number of progenitors *and* lower discrepancy compared to these solutions. Thus, *Carta* provides a systematic approach to evaluate cell differentiation maps with varying number of progenitors and to identify an optimal solution with *k*^*^ progenitors – leading to accurate inference of the cell differentiation map (Methods Section 4.2).

### 2.2 Simulated data

We compared *Carta* to ICE-FASE [47] and *evolutionary coupling* (EvoC) [39], two methods previously used for cell differentiation map inference in lineage tracing studies, on simulated data. ICE-FASE and EvoC use distance-based heuristics calculated from cell lineage trees to perform hierarchical clustering of the cell types to produce the cell differentiation map. Importantly, both of these methods infer cell differentiation maps that are binary trees with *k* = |*S*| − 1 progenitor cell types, where *S* is the set of observed cell types. While *Carta* relies only on the topology of the input cell lineage trees, both ICE-FASE and EvoC additionally require timed cell lineage trees as input.

We simulated two types of cell differentiation maps: (i) trees (not necessarily binary) and (ii) directed acyclic graphs (DAGs). In each case, we generated cell differentiation maps by randomly sampling *k* = 2, 4, 6 progenitor cell types that lead to the generation of *r* = 4, 6 observed cell types in the simulated data (Methods Sections 4.3.1). For each cell differentiation map, we simulated timed cell lineage trees using the simulator in the Cassiopeia [40] platform which employs a generalized Birth-Death model [50] (Methods Section 4.3.2). We then simulated cell type labelings on the vertices of each cell lineage tree, only allowing transitions that exist in the the cell differentiation map. We sampled 50, 100, or 150 cells of each observed cell type from a larger tree such that the total number of sampled cells is 20% of the number of leaves in the larger tree. We provided this pruned cell lineage tree as input to the three methods, and we additionally provided the number *k* of progenitors to *Carta*. This sampling procedure mimics limitations in current lineage tracing technologies in which only a small fraction of cells of the developmental system are sampled for sequencing.

We evaluated the performance of each method by comparing the set 𝒫^*^ of progenitors present in the ground-truth cell differentiation map and the set 𝒫 of progenitors present in the inferred cell differentiation map using two metrics: the Jaccard distance *d*_*J*_ (𝒫, 𝒫^*^) [51] and the *normalized minimum Hamming distance d*_*H*_ (𝒫,𝒫^*^) (Methods Sections 4.3.3). The Jaccard distance evaluates how well 𝒫^*^ matches 𝒫 while the normalized minimum Hamming distance evaluates the deviation of each progenitor in 𝒫^*^ from each progenitor in 𝒫. Both metrics are 0 when the set 𝒫 of inferred progenitors exactly matches the set 𝒫^*^ of ground-truth progenitors.

*Carta*-tree and *Carta*-DAG both outperform ICE-FASE and EvoC when the ground-truth cell differentiation map is a tree (Figure 2a) or DAG (Figure 2b) on all simulation parameters. In the cases where the cell differentiation map is a tree, i.e. Case (i), both *Carta*-tree and *Carta*-DAG either outperform or match the performance of existing methods. Specifically, *Carta*-tree has the lowest Jaccard distance (median 0, mean 0.0119) (Figure 2b), but *Carta*-DAG (median 0, mean 0.0399) also has lower distance than ICE-FASE (0, 0.135) and EvoC (0.4, 0.374). This indicates the ability of *Carta*-DAG to accurately infer cell differentiation maps that are trees even when its output is not restricted to be a tree. Additionally, both *Carta*-tree and *Carta*-DAG as well as ICE-FASE have perfect precision and recall of the ground-truth progenitors in almost all instances (mean precision, mean recall; *Carta*-DAG: 0.975, 0.975, *Carta*-tree: 0.993, 0.993, ICE-FASE: 0.866, 0.999). Similar trends are observed for the normalized minimum Hamming distance metric (Figure 8). In the DAG case, i.e. Case (ii), with 150 sampled cells for each cell type, *Carta*-DAG achieves the lowest Jaccard distance (median 0) (Figure 2a) and normalized minimum Hamming distance (median 0) compared to ICE-FASE (median 0.571 and 0.0365) and EvoC (median 0.67 and 0.0556). Further, *Carta*-tree achieves lower Jaccard distance (median 0.5) and normalized minimum Hamming distance (median 0.0352) than ICE-FASE and EvoC. This indicates that *Carta* is the most accurate tree-restricted method in inferring ground-truth cell differentiation maps that are not restricted to be trees.

**Figure 2:**
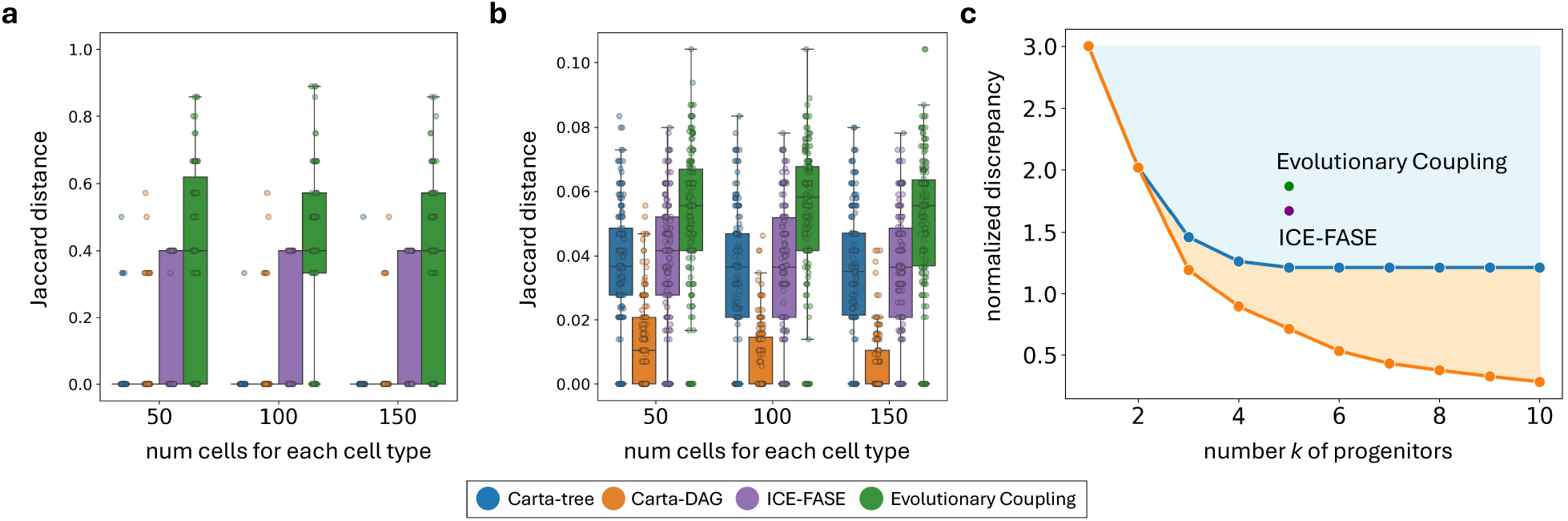
*Carta* outperforms existing methods in inferring cell differentiation maps on simulated data. (a, b) Jaccard distance between the progenitors inferred by each method to the ground-truth progenitors when the cell differentiation map is (a) a tree and (b) a DAG. Box plots show the median and the interquartile range (IQR), and the whiskers denote the lowest and highest values within 1.5 times the IQR from the first and third quartiles, respectively. (c) Normalized discrepancy 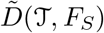(𝒯, *F*_*S*_) of cell differentiation maps, inferred by *Carta*-tree (blue) and *Carta*-DAG (orange), for *k* = 1, … 10 reveals the Pareto fronts; here shown for a simulated cell differentiation map with 6 progenitor cell types, 6 observed cell types, and 150 cells of each observed cell type. Discrepancy of cell differentiation maps inferred by ICE-FASE (purple) and Evolutionary Coupling (EvoC) (green) for their fixed number of 5 progenitors.

*Carta* also defines the Pareto fronts illustrating the trade-off between the discrepancy and the number of progenitors for both tree and DAG cell differentiation maps for varying number *k* of progenitors (Figure 2c). These results show that *Carta*-tree will always infer cell differentiation maps with lower discrepancy than tree-restricted methods such as EvoC and ICE-FASE, and that less-constrained *Carta*-DAG will always infer cell differentiation maps with lower discrepancy than any tree restricted method. Further, while EvoC and ICE-FASE are restricted to only infer cell differentiation maps with |*S*| − 1 progenitors, *Carta*-tree can infer differentiation maps with a wide range of number of progenitors. Note that here and below, we report the *normalized discrepancy* 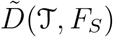(𝒯, *F*_*S*_), which is the discrepancy divided by the total number of ancestral cells across the input lineage trees.

### 2.3 Cell differentiation mapping of Trunk-Like Structures (TLS) – an *in vitro* model of the mammalian embrynonic trunk

We compared *Carta* and several other methods in inferring the routes of differentiation during mammalian trunk development. Specifically, we applied *Carta*, Fitch, PhyloVelo [52], ICE-FASE [47] and Evolutionary Coupling [38] (Methods Section 4.5) to cell lineage trees derived from single-cell CRISPR-Cas9-based lineage tracing of an *in vitro* embryoid model called Trunk-Like Structures (TLS) [29]. TLS mirrors post-occipital mammalian trunk development and is particularly suited for studying the differentiation dynamics of neuromesodermal progenitor (NMP) cells. NMPs are a pool of self-renewing progenitors that differentiate into both the neural tube, which forms the future spinal cord, and the flanking somitic mesoderm, which form future vertebrae and muscle cells of the trunk (Figure 3a) [48]. Given their bipotent nature, NMPs are particularly interesting as they produce cells of two germ layers in the posterior embryo, the neuroectoderm and the paraxial mesoderm, that are classically considered to come from separate origins [53–55]. This dataset consists of 14 cell lineage trees with a total of 6570 cells labeled by 6 observed cell types derived from the gene expression measurements: Endoderm (233 cells), Endothelial (124 cells), Primordial germ cell-like cells (PGCLCs; 233 cells), Somites (3188 cells), Neural Tube (2289 cells), and NMP (513 cells) (see Methods Section 4.4.1 for details).

**Figure 3:**
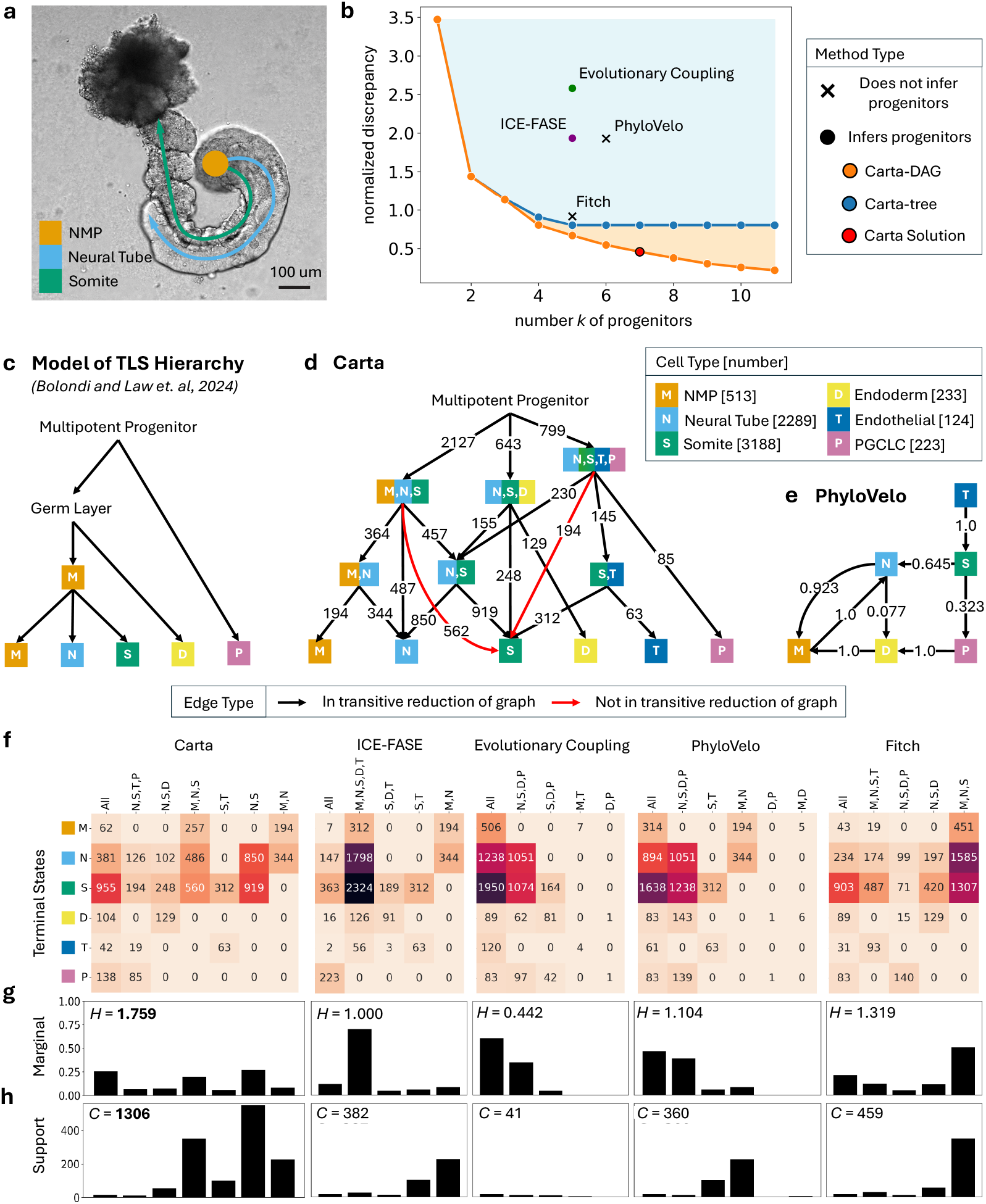
*Carta* accurately infers the cell differentiation map of Trunk-Like Structures (TLS), an *in vitro* model of mammalian trunk development. (a) A representative live-imaged TLS structure at 120h with NMP progenitor pool (orange) and elongating neural tube (light blue) and somite (green) structures. (b) Normalized discrepancy 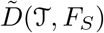(𝒯, *F*_*S*_) of cell differentiation maps, inferred by *Carta*-tree (blue) and *Carta*-DAG (orange), for increasing number *k* of progenitors revealing the Pareto fronts. Discrepancy of cell differentiation maps inferred by existing methods that infer unobserved progenitors (·) and do not infer unobserved progenitors (X) are also shown. (c) Canonical model of TLS differentiation [29]. (d) Cell differentiation map inferred by *Carta*-DAG, where edges are annotated by number of cells that traverse the cell type transition. Legend indicates the number of cells of each cell type. (e) Cell differentiation map inferred by PhyloVelo. Weight of each edge is the inferred transition probability between two cell types. (f) The number of cells that directly transition from progenitor cell types (rows) to observed cell types (columns) for the cell differentiation maps inferred by each method. (g) The marginal distribution, or the proportion of total cells that derive from each progenitor, and the corresponding entropy *H* of each distribution. (h) The support, or the number of internal nodes in the lineage trees with the exact potency as each progenitor.

We compared the cell differentiation maps generated by *Carta* with varying number of progenitors to the differentiation maps inferred by existing methods. Both modes of *Carta*, i.e. *Carta*-tree and *Carta*-DAG, consistently infer cell differentiation maps with lower discrepancy compared to existing methods for the same number of progenitors (Figure 3b). For example, the normalized discrepancies of cell differentiation maps with *k* = 5 progenitors that *Carta*-tree and *Carta*-DAG infer are 0.802 and 0.668, respectively In contrast, ICE-FASE, EvoC and Fitch infer cell differentiation maps that have 5 progenitors and normalized discrepancy of 1.936, 2.580 and 0.915, respectively (see Methods Section 4.8 for details). PhyloVelo infers a map with 6 progenitors with normalized discrepancy of 1.930 compared to 0.802 and 0.546 for *Carta*-tree and *Carta*-DAG respectively with *k* = 6. We determined the optimal number *k*^*^ = 7 progenitors in the cell differentiation maps (normalized discrepancy 0.458) by identifying the elbow in the Pareto fronts derived using *Carta*-DAG (See Methods Section 4.6 for details).

The cell differentiation map inferred by *Carta* (Figure 3d) agrees with known features of trunk developmental progression. *Carta*-tree infers a cell differentiation map in which the relative ordering of commitment of observed cell types is – PGCLC, endoderm, endothelial, NMP, somites and neural tube (Supplementary Figure 6). This is consistent with the canonical model of TLS differentiation in which the fate of PGCLC and endoderm cells is committed earlier compared to the NMP, somite and neural tube cells (Figure 3c) [48]. This is also reflected in the *Carta*-DAG cell differentiation map, in which endothelial, endoderm, and PGCLC cells derive from progenitors with larger potencies (mean progenitor potency size: 4.5, 4.0 and 5.0, respectively) compared to the more closely related NMP, somitic, and neural tube cells which arise from progenitors with more restrictive potencies (mean progenitor potency size: 3.7, 3.3 and 3.3, respectively).

A key insight of the *Carta*-DAG cell differentiation map is the convergent differentiation of somite cells, with one origin stemming from shared ancestry with neural tube cells and an alternate origin indicating shared ancestry with endothelial cells via the presence of the {endothelial, somite} progenitor. This is consistent with previous *in vivo* studies that have found evidence for a secondary pathway towards the production of the trunk endothelium [56, 57]. Such instances of convergent differentiation cannot be revealed by methods such as ICE-FASE and EvoC that infer only tree-structured cell differentiation maps in which each cell type arises from a single developmental trajectory.

*Carta* further reveals the progenitor dynamics as well as the commitment bias of NMPs, i.e. the proportion of NMPs committing to each downstream state. The *Carta*-DAG differentiation map includes NMPs in multiple known stages of development [53–55]. The {NMP} cell type represents observed undifferentiated NMPs, the {neural tube, somite} cell type represent ancestral NMPs that existed in the past, and the {NMP, neural tube, somite} cell type represent NMP cells that both self renew and are differentiating. Further, the {NMP, neural tube} cell type represents NMP cells that are only observed differentiating into neural tube cells, and the NMP, somite cell type represents NMP cells that are only observed differentiating into somitic cells. Notably, all of the different instances of progenitor cell types such as {NMP, neural tube} and neural tube, somite can only be represented simultaneously in a DAG structure and not a tree structure. We observe that the *Carta*-DAG cell differentiation map only includes the {NMP, neural tube} and not the {NMP, somite} progenitor, suggesting that NMP cells in this system have a higher propensity to commit to a neural rather than somitic fate (Figure 3c). This bias towards neural fate supports previous analyses that NMP cells gradually shift their differentiation potential towards the neural fate during TLS development [29].

In contrast, methods where all progenitor cell types are assumed to be observed – such as Fitch and PhyloVelo – infer cell differentiation maps that are not well supported by the literature. Many spurious cell type transitions exist in the differentiation map produced by PhyloVelo (Figure 3d). For example, somitic cells differentiate into PGCLC, endoderm, and neural tube cells. Further, endothelial cells differentiate into somites and endoderm cells differentiate into NMPs. In these instances, observed cell types are shown to transition directly to each other when it is known that these cell types are related through progenitor cell types that are potent for each of them. This highlights that the deficiencies of the assumption that all progenitors are observed. Additionally, PhyloVelo does not correctly infer the hierarchical differentiation process as the differentiation map shows neural tube cells can differentiate back to the NMP state (Figure 3d). The cell differentiation maps inferred by ICE-FASE, EvoC, and Fitch also have poor agreement with the reported developmental routes in TLS (Supplementary Figure 6).

The progenitors inferred by *Carta* are better supported by the cell types of descendants of ancestral cells in the cell lineage trees. We demonstrate this advantage using two metrics. First, we calculate the distribution of the number of cells of each observed cell type that directly arise from the progenitor cell type inferred by each method (Figure 3f, Methods Section 4.8). The progenitors inferred by *Carta* produce a more uniform distribution Figure 3g) quantified by the higher entropy (*H* = 1.759) of the distribution compared to existing methods (ICE-FASE: *H* = 1.0, EvoC: *H* = 0.820, PhyloVelo: *H* = 1.104, Fitch: *H* = 1.319). Moreover, for ICE-FASE, EvoC and PhyloVelo, the proportion of cells arising from the two progenitors that account for the most cells (ICE-FASE: 0.814, EvoC: 0.954, PhyloVelo: 0.859, Fitch: 0.719) is substantially larger than the proportion of 0.525 for *Carta*. Second, we calculate the *support* of the inferred progenitors, i.e. the number of ancestral cells where the set of cell types of the descendants exactly match the potencies of the inferred progenitors (Figure 3h). Progenitors inferred by *Carta* have much higher support *C* = 1306 compared to existing methods (ICE-FASE: *C* = 382, EvoC: *C* = 41, PhyloVelo: *C* = 360, Fitch: *C* = 459), indicating that *Carta* differentiation map provides a better fit with the input cell lineage trees.

### 2.4 Carta reveals the hierarchy of progenitors during mouse hematopoiesis

We applied *Carta* and several existing methods to a single-cell lineage tracing dataset of mouse hematopoiesis [30] and compared the resulting cell differentiation maps. This dataset was obtained by inserting random and heritable lentiviral barcodes in mouse hematopoietic stem cells (HSCs), which were then allowed to differentiate *in vitro*, with the culture sampled at days 2, 4 and 6. Single-cell RNA sequencing was performed to simultaneously measure the barcodes and gene expression of the 11778 sampled cells. The barcode measurements were used to construct 5624 star-shaped cell lineage trees, one for each unique barcode shared across multiple cells (Methods Section 4.4.2). The cells were annotated into 9 observed cell types based on gene expression – Megakaryoctyes (Meg), Erythrocytes (Ery), Mast cells (Ma), Basophils (Ba), Eosinophil (Eo), Neutrophils (Neu), Monocytes (Mo), Dendritic cells (DC) and Lymphoid (L), with the remaining 22387 cells marked as undifferentiated (Figure 4a).

**Figure 4:**
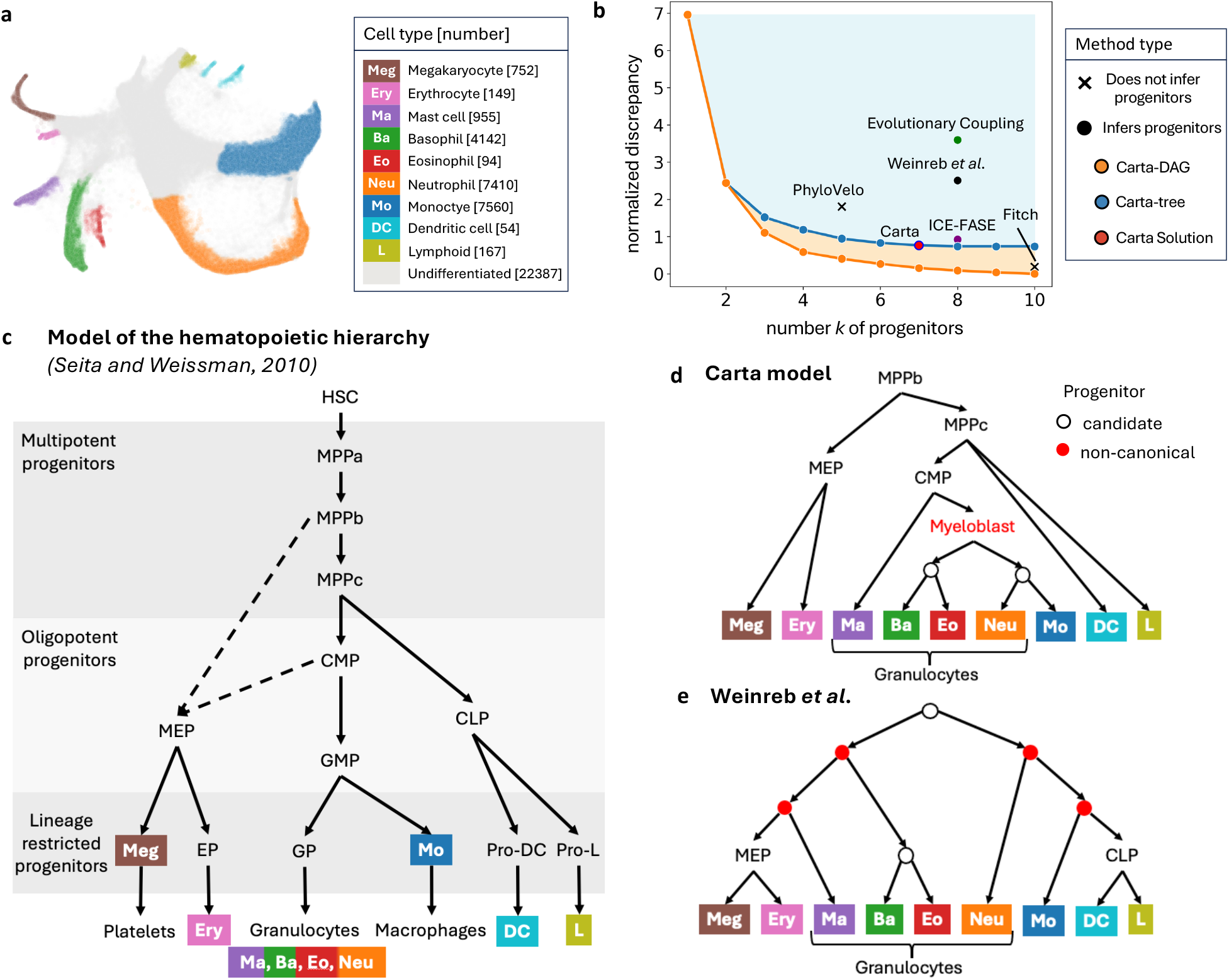
*Carta* recapitulates canonical model of mouse hematopoiesis from lentiviral barcoding-based lineage tracing data. (a) Low-dimensional visualization [58] of scRNA-seq of 43670 clonally barcoded cells in varying stages of mouse hematopoesis differentiation [30]. Cells are colored by cell type, and legend contains the number of cells of each cell type. (b) Normalized discrepancy 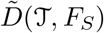(𝒯, *F*_*S*_) of cell differentiation maps inferred by *Carta* and existing methods with varying number of progenitors. (c) Canonical model of the hierarchy of progenitors during mouse hematopoiesis from [59]. Dashed arrows show alternate routes of differentiation that have been suggested in previous studies. (d) Cell differentiation map inferred by *Carta* and (e) distance-based heuristic employed by Weinreb *et al*. [30]. Red indicates inferred progenitors that are non-canonical, i.e. do not agree with the canonical model shown in (c).

**Figure 5:**
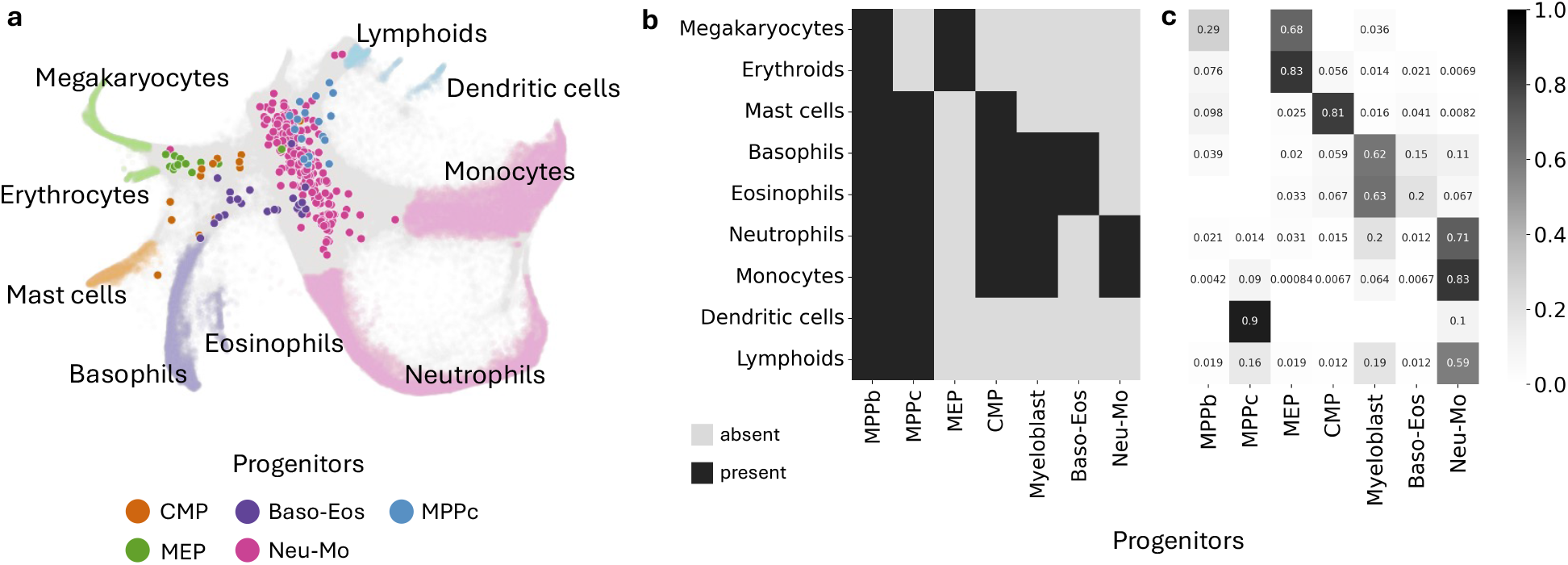
*Carta* predicts the fate of undifferentiated hematopoietic progenitor cells. (a) Progenitor predictions given by *Carta* (colored dots) for undifferentiated cells sampled at day 2. (b) Potencies of the inferred progenitor cell types. (c) The proportion of undifferentiated cells (from day 2, 4, and 6) that are closest in transcriptional space to the indicated observed cell type for each predicted progenitor.

We compared the differentiation maps inferred by both modes of *Carta, Carta*-tree and *Carta*-DAG, to cell differentiation maps published in the original study (Weinreb *et al*. [30]) and inferred using existing methods – Fitch, PhyloVelo [52], ICE-FASE [47], and Evolutionary Coupling [39] (Figure 4b) (Methods Section 4.5). *Carta*-tree and *Carta*-DAG both infer solutions with *k*^*^ = 7 progenitors and normalized discrepancy of 0.154 and 0.762, respectively. PhyloVelo infers a cell differentiation map with only 5 progenitors, but much higher normalized discrepancy 1.809, while Fitch infers a map with 9 progenitors with low normalized discrepancy of 0.186. ICE-FASE, EvoC, and Weinreb *et al*. [30] infer tree-structured cell differentiation maps comprising of 8 progenitors, but with higher normalized discrepancy of 0.921, 3.598, 2.51, respectively compared to *Carta*-tree with the same number (*k* = 8) of progenitors and normalized discrepancy = 0.738. Since the canonical model of hematopoiesis [59] is also tree-structured, we focus our attention on the optimal solution inferred by *Carta*-tree with *k*^*^ = 7 progenitors (Methods Section 4.6).

The cell differentiation map inferred by *Carta* aligns more closely with the canonical model of hematopoiesis [59] (Figure 4c) compared to the hierarchy of progenitors published in the original study [30]. *Carta* infers that the myeloid cells (Ma, Ba, Eo, Neu and Mo) originate from a common unobserved progenitor cell type, which we identify as the common myeloid progenitor (CMP) that is consistent with the canonical model of hematopoiesis [59–61] (Figure 4d). Additionally, *Carta* also infers an intermediate non-canonical progenitor, which we identify as myeloblast [62], with potency for Ba, Eo, Neu and Mo cells. In contrast, Weinreb *et al*. [30] suggest that the myeloid cells separate into two trajectories (first containing Ma, Ba, Eo and second containing Neu and Mo) very early during differentiation when the cells are still multipotent progenitors (Figure 4e). Additionally, *Carta* identifies an unobserved progenitor restricted to megakaryocytes and eythrocytes, known as the megakaryocyte-erythrocyte progenitor (MEP), which arises directly from multipotent progenitor (MPP) cells. This finding is consistent with previous studies that have found evidence that in mouse, MPP give rise to MEP without passing through the CMP [59, 63–66]. While Weinreb *et al*. [30] also identify MEP, they propose that it originates from a non-canonical progenitor that is potent for megakaryocytes, eythrocytes and mast cells. *Carta* also correctly infers that lymphoid and dendritic cells belong to a differentiation trajectory that separates early from the other cell types (myeloids, megakaryocytes and erythrocytes) during hematopoiesis [59, 64]. However, *Carta* is not able to identify the presence of the common lymphoid progenitor (CLP), possibly due to low sampling of lymphoid and dendritic cells in the data (18 and 22 cells, respectively). In contrast, Weinreb *et al*. [30] identify the CLP, but suggests that it originates from a non-canonical hierarchy of progenitors with potency for Neutrophils and Monocytes. Finally, the *Carta* cell differentiation tree has the lowest Robinson-Foulds distance [67] with the canonical tree (1; maximum possible 7), compared to the tree inferred by Weinreb *et. al* (4; maximum possible 8), the ICE-FASE tree (6; maximum possible 8), and the EvoC tree (2; maximum possible 8) (Supplementary Figure 7).

**Figure 6:**
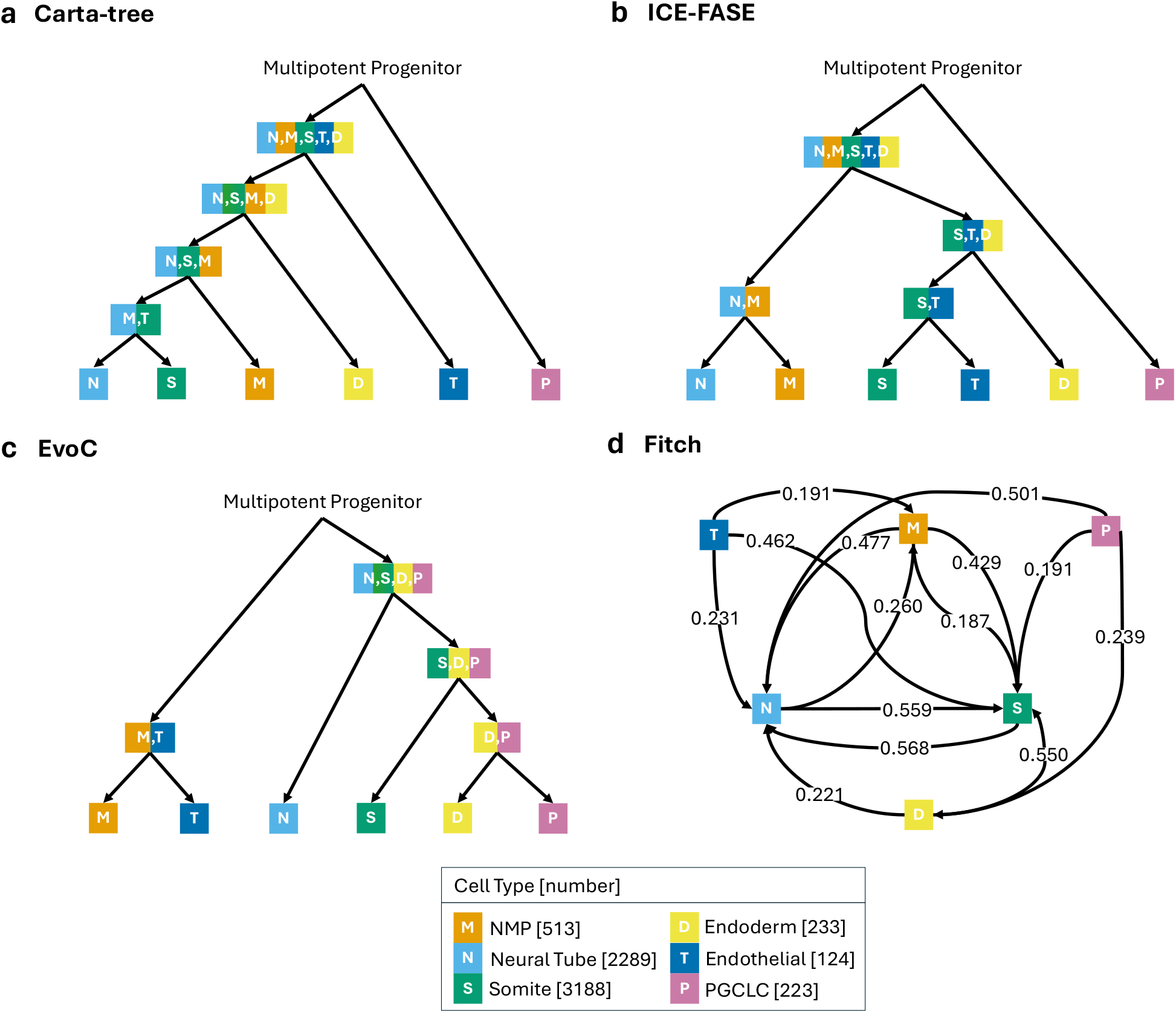
Graphical representations of cell differentiation maps inferred for the TLS dataset by other methods. (a) *Carta*-tree cell differentiation map. (b) ICE-FASE cell differentiation map. (c) EvoC cell differentiation map. (d) Fitch cell differentiation map. Edge weights indicate the normalized transition frequency from one observed cell type to another. Only edges with frequency 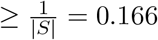 are shown.

**Figure 7:**
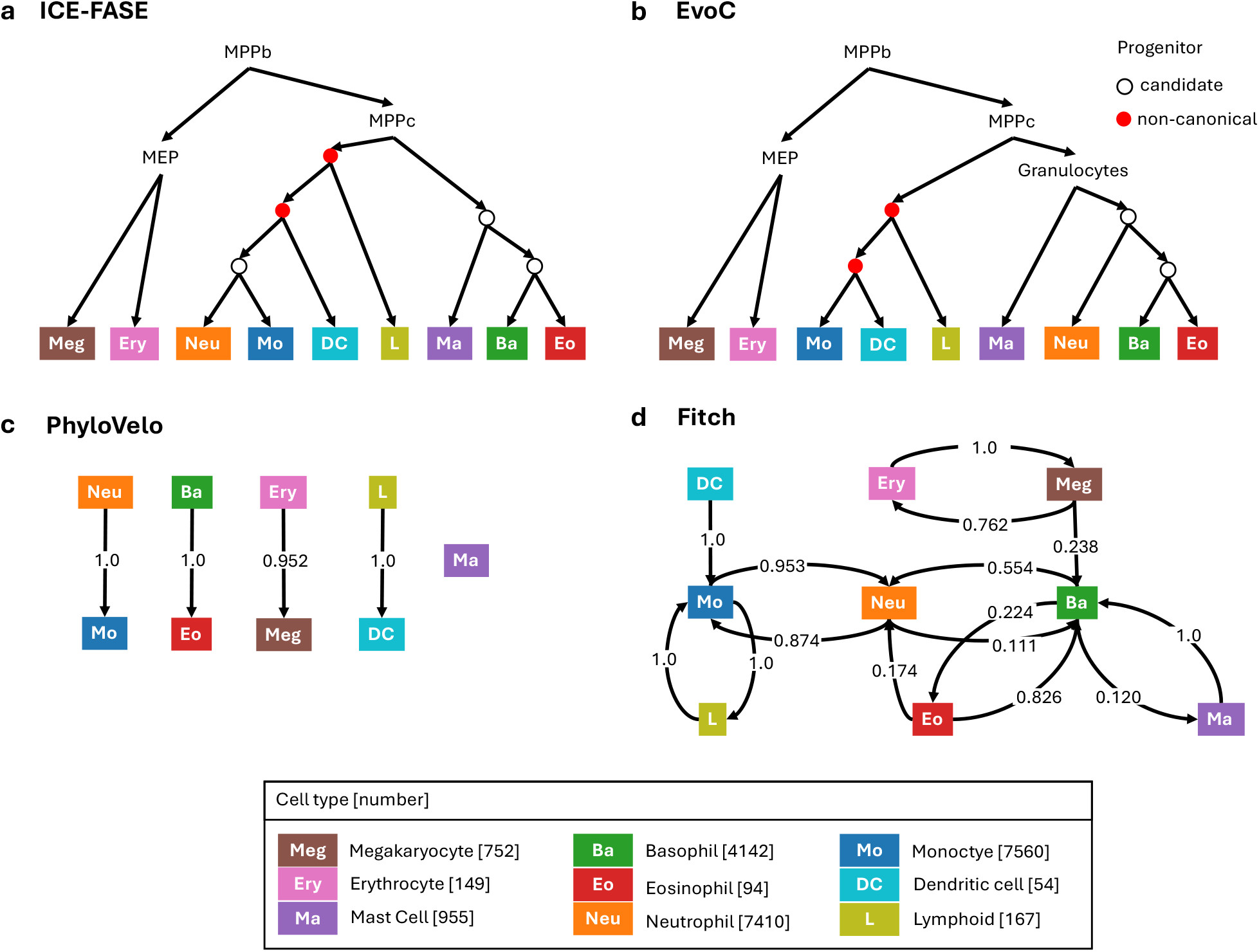
Graphical representations of cell differentiation maps inferred for the dataset from Weinreb *et. al* by other methods. (a) ICE-FASE cell differentiation map. Red indicates inferred progenitors that are non-canonical, i.e. do not agree with the canonical model. (b) EvoC cell differentiation map. Red indicates inferred progenitors that are non-canonical, i.e. do not agree with the canonical model. (c) PhyloVelo cell differentiation map. Edge weights indicate the normalized transition frequency from one observed cell type to another. Only edges with frequency 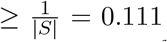 are shown. (d) Fitch cell differentiation map. Edge weights indicate the normalized transition frequency from one observed cell type to another. Only edges with frequency 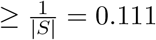 are shown.

**Figure 8:**
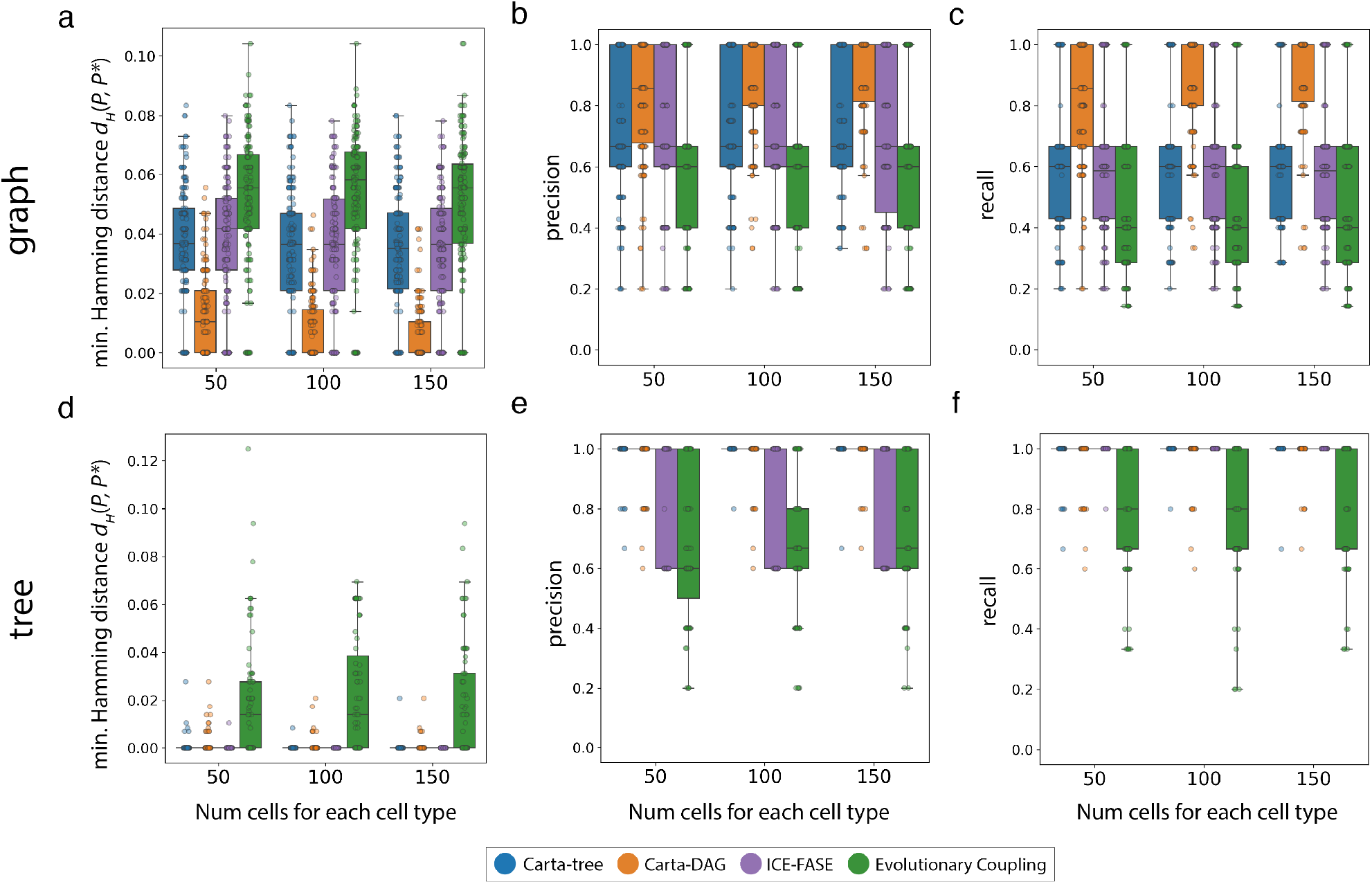
Additional metrics for simulated data. (a, d) Normalized minimum Hamming distance between the progenitors inferred by each method to the ground-truth progenitors when the cell differentiation map is (a) a DAG and (d) a tree. Box plots show the median and the interquartile range (IQR), and the whiskers denote the lowest and highest values within 1.5 times the IQR from the first and third quartiles, respectively. (b, e) Precision (the proportion of inferred progenitors that are in the ground truth cell differentiation map) for each method when the cell differentiation map is (b) a DAG and (e) a tree. (c, f) Recall (the proportion of progenitors in the ground truth cell differentiation map that are in the inferred set of progenitors) for each method when the cell differentiation map is (c) a DAG and (f) a tree.

We examine the concordance between the progenitor cell types of undifferentiated cells predicted by *Carta* and the gene expression of these cells. Specifically, we defined the progenitor cell type of undifferentiated cells sampled at day 2 based on the progenitor cell type inferred by *Carta* for their ancestors in the cell lineage trees. We find that the undifferentiated cells have similar gene expression to the observed cell types in the potency set; i.e. the cell types that *Carta* predicts the undifferentiated cell will differentiate into (Figure 5a). We quantify this similarity by comparing the predicted fate of undifferentiated cells to the cell type cluster of the closest cell in gene expression space (Methods Section 4.9). We observe a high degree of overlap between predicted potency and closest mature cell type in the cases where the inferred progenitor is potent for that cell type (Figure 5b-c). Since *Carta* uses only lineage information and not gene expression in inferring progenitors, these results provide orthogonal validation for the progenitor cell types inferred by *Carta*.

## 3 Discussion

We introduce *Carta*, an algorithm to infer cell differentiation maps from cell lineage trees while accounting for limitations in high-throughput lineage tracing data such as limited sampling of cells. *Carta* employs a new mathematical model of differentiation maps, in which progenitor cell types are defined by their potency, i.e. the set of cell types that can be attained by their descendants. This model allows for the inference of transient progenitor cell types that arise during development but may not be observed in the lineage tracing data. A key insight of our work is that there exists a trade-off between the number of progenitors in the cell differentiation map (a measure of the *complexity* of the map) and how well the map fits the input cell lineage trees (*discrepancy*). *Carta* explicitly evaluates this trade-off by deriving the Pareto front of cell differentiation maps and selecting a map with an optimal number of progenitor cell types.

We demonstrate the advantages of *Carta* compared to other methods on simulated and real data from multiple single-cell lineage tracing technologies. On simulated lineage tracing data, *Carta* reconstructs more accurate cell differentiation maps compared to existing methods under varying simulation parameters. On CRISPR-Cas9-based lineage tracing data of Trunk-like Structures, an *in vitro* model of mammalian trunk development, *Carta* provides insights into differentiation of neuro-mesodermal progenitors (NMPs) and convergent differentiation of somites, features that are not revealed by the restrictive frameworks of existing methods. Additionally, on lentiviral barcoding-based lineage tracing data of mouse hematopoiesis, the cell differentiation map that *Carta* infers has better agreement with canonical model of mouse hematopoiesis compared to maps inferred by existing methods and has high concordance with the gene expression measurements.

There are several limitations of *Carta* which present opportunities for future development. First, *Carta* takes cell lineage trees derived from lineage tracing data as input, but these trees are not always accurate [44]. Joint inference of cell lineage trees and a cell differentiation map might lead to improvement in the accuracy of both the trees and the differentiation map. Second, we defined the discrepancy by counting unsampled descendant resulting in a maximum parsimony framework to infer cell differentiation maps. A promising direction for future research is derivation of a maximum likelihood-based framework that employs a probabilistic model for cell differentiation and fate commitment during development. Third, *Carta* quantifies the complexity of the differentiation map by the number of progenitors, but complexity could also be described in terms of the number and type of transitions (See Supplementary Section A.1). Finally, *Carta* assumes that progenitors do not regain potency for a cell type once it is lost during differentiation, and thus does not model dedifferentiation. While this assumption is reasonable for most normal developmental systems, it is not hold in aberrant systems such as cancer. Indeed, dedifferentiation has been recognised as a major mechanism of cancer progression, cancer cell plasticity and immune evasion [68–71]. Extending *Carta* to allow dedifferentiation would enable further application to study cancer development.

Finally, investigations of developmental systems are increasingly utilizing varied high-throughput technologies including spatial RNA sequencing [31, 72, 73] and single-cell multi-modal sequencing [74]. Combining lineage tracing with multi-modal single-cell and spatial sequencing is crucial for measuring the interplay between microenvironment, epigenetic regulation and lineage of the cells. We envision that *Carta* will play a crucial role in distinguishing the relative contributions of cell lineage, cell differentiation, and spatial location during development and provide a foundation for future development of algorithms for cell differentiation mapping.

## 4 Methods

### 4.1 Definition and inference of cell differentiation maps

A *cell differentiation map F*_*S*_ describes the differentiation of cells into observed cell types *S*. Here, we give a formal definition of a cell differentiation map *F*_*S*_ and formulate the problem of inferring a cell differentiation map from a set 𝒯 of cell lineage trees (Problem 4.2).

We define a *cell differentiation map F*_*S*_ to be a vertex-labeled directed graph whose sinks – i.e. vertices with outdegree *d* = 0 – are the observed cell types *S*, and whose whose internal vertices – i.e. vertices with outdegree *d* > 0 – are the progenitor cell types. The directed edges of *F*_*S*_ describe the cell type transitions that occurred during development. Each sink vertex (observed cell type) *t* ∈ *S* is labeled by the singleton set {*t*} (or for simplicity by an element of *S*) and each internal vertex (progenitor cell type) is labeled by a potency set, i.e. a subset of *S*.

We model development as a process in which cells progressively lose potency and do not regain potency for a cell type once it is lost. Thus, *F*_*S*_ is a directed acyclic graph – i.e. does not have directed cycles – in which the root of *F*_*S*_ has label *S* indicating the totipotent cell with potency *S*, and the internal vertices have unique labels that satisfy the following two conditions. First, since we assume cells only lose potency during development, every directed edge (*P, P* ^*′*^) in *F*_*S*_ satisfies *P* ^*′*^ ⊂ *P*. Second, by definition of potency, for each cell type *t* ∈ *S* there exists a directed path in *F*_*S*_ from a progenitor *P* to a observed cell type {*t*} if and only if it is potent for the cell type *t*, i.e. *t* ∈ *P*. Consequentially, the vertex set P_*S*_ of a cell differentiation map *F*_*S*_ always contains the totipotent cell *S*, and the singleton set {*t*} for each observed cell type *t* ∈ *S*.

The cell types of ancestral cells are determined by a labeling of the internal vertices of 𝒯 (ancestral cells) by the vertices of the cell differentiation map *F*_*S*_ (cell types). Such a labeling must be *compatible* with the trees 𝒯 and cell differentiation map *F*_*S*_, i.e. it must satisfy the following two conditions. First, each ancestral cell in a cell lineage tree must be labeled by a potency that contains all the observed cell types of its descendants in the tree. Second, cell type transitions determined by the labeling – i.e. edges in the lineage trees connecting vertices labeled by distinct cell types – must be supported by the cell differentiation map *F*_*S*_. More formally, for every edge (*u, v*) in a cell lineage tree, there must exist a path from ℓ(*u*) to ℓ(*v*) in *F*_*S*_.

For cell lineage trees 𝒯 and a cell differentiation map *F*_*S*_, there may be multiple compatible labelings. We evaluate a labeling ℓ by its discrepancy, defined as the number of instances when a cell type in the potency ℓ(*v*) of an ancestral cell *v* is not observed in its descendants, i.e. the leaves of the subtree rooted at *v*. More formally,

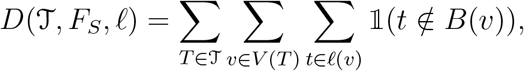

where 𝟙 is the indicator function and *B*(*v*) is the set of observed cell types of the descendants of cell *v*.

We define the discrepancy between the cell lineage tree 𝒯 and a cell differentiation map *F*_*S*_ by the minimum discrepancy obtained over all compatible labelings, i.e.

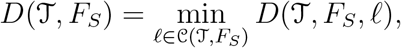

where 𝒞(𝒯, *F*_*S*_) is the set of compatible labelings for cell lineage trees 𝒯 and cell differentiation map *F*_*S*_. A more general description of discrepancy is given in Supplementary Section A.1.

As such, evaluating the discrepancy of a given cell differentiation map *F*_*S*_ with a set of cell lineage trees 𝒯 is equivalent to finding a compatible vertex labeling ℓ that minimizes the induced discrepancy *D*(𝒯, *F*_*S*_, ℓ). We refer to this as the Progenitor Labeling Problem (PLP) and formally pose it as follows.

#### Problem 4.1

(Progenitor Labeling Problem (PLP)). *Given a set* 𝒯 *of cell lineage trees and cell differentiation map F*_*S*_, *find a valid labeling* ℓ *that minimizes the discrepancy D*(𝒯, *F*_*S*_, ℓ).

This is an analog of the *small parsimony* problem [75], and we show that it can be solved by a dynamic program by adapting the Sankoff’s algorithm [76] (Supplementary Section A.3).

In practice, we only observe the cell lineage trees 𝒯 and must infer the cell differentiation map *F*_*S*_. Due to technical limitations in current lineage tracing technologies, such as limited sampling of cells, inferring a map *F*_*S*_ with the minimum the discrepancy *D*(T, *F*_*S*_) may lead to large number of progenitors, many of which may be false positives. As such, we pose the Cell Differentiation Map Inference Problem (CDMIP) of inferring a cell differentiation map with minimum discrepancy for a fixed number *k* of progenitors.

#### Problem 4.2

(Cell Differentiation Map Inference Problem (CDMIP)). *Given cell lineage trees* 𝒯 *with observed cell types S, and integer k, find a cell differentiation map F*_*S*_ *with k progenitors such that D*(T, *F*_*S*_) *is minimized*.

An interesting special case of the CDMIP problem is when the differentiation map is restricted to be a tree. We define this problem as follows.

#### Problem 4.3

(Cell Differentiation Tree Inference Problem (CDTIP)). *Given cell lineage trees* 𝒯 *with observed cell types S, and integer k, find a cell differentiation tree F*_*S*_ *with k progenitors such that D*(𝒯, *F*_*S*_) *is minimized*.

We show that both the CDMIP and CDTIP problems are NP-complete (see Supplementary Section A.5, proofs in Supplementary Section A.6).

### 4.2 *Carta*: an algorithm for cell differentiation mapping

We develop *Carta*, an algorithm, to infer a cell differentiation map *F*_*S*_ from cell lineage trees 𝒯 that balances the trade-off between the discrepancy *D*(T, *F*_*S*_) and the number *k* of progenitors in the cell differentiation maps. *Carta* allows inference of DAG and tree-structured cell differentiation maps by solving this multi-objective optimization problem in two steps, which we detail below.

First, *Carta* finds the cell differentiation map with minimum discrepancy for each number *k* of progenitors across a range of values of *k*. This reveals the Pareto front indicating the minimum discrepancy obtained over differentiation maps for each fixed number *k* of progenitors. *Carta* has two modes, *Carta*-DAG for DAG-structured cell differentiation maps and *Carta*-tree tree-structured cell differentiation maps. For a fixed number *k* of progenitors, *Carta*-DAG and *Carta*-tree use on mixed integer linear programs (MILPs) to solve the Cell Differentiation Map Inference Problem (CDMIP, Problem 4.2) and the Cell Differentiation Tree Inference Problem (CDTIP, Problem 4.3), respectively. The MILPs are solved using the Gurobi optimizer [77] in Python and the details of the MILP formulations are described in the Supplementary Section A.2.

Second, *Carta* determines the optimal number *k*^*^ of progenitors by identifying the *elbow* of the Pareto front. To this end, we use *kneedle* [78], a heuristic algorithm that finds the point of maximum curvature on the Pareto front (details in Methods Section 4.6). The edges of the cell differentiation map are determined by including all cell type transitions that appear frequently in the labeled cell lineage trees (details in Methods Section 4.7).

### 4.3 Simulation details

#### 4.3.1 Simulating DAG cell differentiation maps

As described in 2.2, we generated two sets of simulations, one where the ground truth cell differentiation map is a DAG and the other where it is a tree. We generated a random DAG-structure cell differentiation map as follows. For a fixed number *k* of progenitors and set *S* of observed cell types, we randomly sampled *k* progenitors from the power set of *S*. These progenitors, along with each observed cell type *t* ∈ *S* and *S* (the totipotent root progenitor), form the vertices in the cell differentiation map. We defined the edges by first build a graph with a directed edge (*P, P* ^*′*^) in the graph if *P* ⊆ *P* ^*′*^ and then taking the transitive reduction of this graph.

We generated a random tree-structured cell differentiation maps with *k* internal vertices (progenitors) and *S* observed cell types using the following iterative process. We initialized the cell differentiation tree as a single vertex. At each iteration, we added a child to an existing vertex chosen uniformly at random. We terminated the process when the map had exactly *k* internal vertices and |*S*| leaves after collapsing unifurcations. The cell types are assigned to these leaves with a one-to-one mapping uniformly at random.

#### 4.3.2 Simulating cell lineage tree from a given cell differentiation map

For each simulated cell differentiation map *F*_*S*_, we simulated time-resolved binary cell lineage trees that follow the differentiation routes specified by that map. To generate tree topologies, we used the generalized forward-time birth-death simulator included in the Cassiopeia platform [40]. Let *z* be the number of cells sampled per extant cell type and let *α* be the subsampling rate. The process terminates when 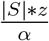 extant tips are sampled. We drew birth waiting times from a shifted exponential distribution with a shift constant of *c* = 0.01, and estimated the birth and death rates to produce trees with total times of around 1 for the given number of extant tips. We then normalized the branch lengths of *T* such that the longest path from the root to one of a leaf of the tree is of length 1 to match the times on *F*_*S*_.

We simulated cell type differentiation in two steps. First, we assigned a differentiation time for each cell type transition in the cell differentiation map. Specifically, we annotated each vertex of the cell differentiation map by a time between 0 and 1 representing the time of arrival of that cell type such that if vertex *u* precedes vertex *v*, then *τ* (*u*) *< τ* (*v*). These times are determined by a process in which we iterated through paths in the cell differentiation map from root to sink, and on each iteration annotated the length of each edge in a path by evenly splitting the remaining length of that path amongst its edges. The time of each vertex is the sum of the path length from the root. Second, we randomly labeled the ancestral cells of each cell lineage tree *T* with cell types such that cell type transitions in *T* are consistent with the cell type transitions in the cell differentiation map *F*_*S*_. To achieve this, we first initialized the label ℓ(*r*(*T*)) of the root vertex *r* of cell lineage tree *T* as the totipotent progenitor *S*. Let *τ*_*T*_ and 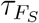 be the timepoint annotation function for the cell lineage tree *T* and cell differentiation map *F*_*S*_, respectively. We performed a depth-first, preorder traversal of the edges (*u, v*) ∈ *E*(*T*) of the lineage tree such that we annotate ℓ(*v*) as ℓ(*u*) if 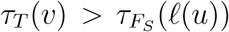 and otherwise a randomly sampled descendant of ℓ(*u*) in the *F*_*S*_. Finally, once each cell in the cell lineage is annotated with a progenitor label, we randomly sampled the specified number *z* = 50, 100, or 150 of cells labeled with each extant cell type in *S*. We took the subtree induced by the sampled cells as well as the cell type labelings of the leaves of this tree as the final inputs to our cell differentiation map inference algorithms (Section 2.2).

#### 4.3.3 Simulation metrics

We evaluate the inferred cell differentiation maps against the simulated ground-truth cell differentiation maps using two metrics that quantify the difference in the progenitors in each:

1. *Jaccard distance d*_*J*_ (𝒫, 𝒫^*^) [51]:

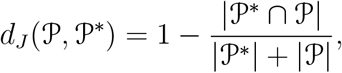

where 𝒫^*^ and 𝒫 are the ground-truth and the inferred set of progenitors, respectively. The Jaccard distance *d*_*J*_ (𝒫, 𝒫^*^) is 0 if and only if the set 𝒫 of inferred progenitors exactly match the set 𝒫^*^ of ground-truth progenitors.
2. The *normalized minimum Hamming distance d*_*H*_ (𝒫, 𝒫^*^):

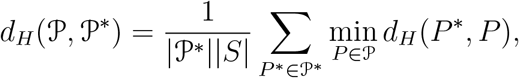

where Hamming distance [79] *d*_*H*_ (*P*^*^, *P*) = |*P* ^*^ *\ P*| + |*P \ P*^*^|.

Intuitively, the Hamming distance of two progenitors is defined as the size of the symmetric difference of the two progenitors and would be 0 if and only if the two progenitors are identical. The normalized minimum Hamming distance is the sum of the minimum Hamming distance between an inferred progenitor and all of the progenitors in the ground-truth, normalized by the number |𝒫^*^ | of ground-truth progenitors and number |*S*| of observed cell types. As such, *d*_*H*_ (𝒫, 𝒫^*^) is 0 if and only if each ground-truth progenitor is present in the inferred set 𝒫 of progenitors.

### 4.4 Data processing details

#### 4.4.1 Processing of TLS data

We obtained 14 cell lineage trees (Table 1) that record the cell division of 14 Trunk-like structures generated in [29] (Section 2.3). These lineages were generated using scRNA-seq readout from mouse embryonic stem cells engineered with CRISPR-Cas9 lineage tracing technology. This scRNA-seq data was then input to the Cassiopeia lineage pre-processing and reconstruction package [40]. The branch lengths are not given by Cassiopeia, and hence we used unit branch lengths. Each observed cell (leaf) in each cell lineage tree was assigned a cell type by a previously published reference [48]. We grouped all somite cell subtypes (Somite (−1), Somite 0, Somite, Sclerotome-like, and Dermomyotome-like) into one umbrella type “somite”, and we grouped NeuralTube1 and NeuralTube2 cell types into one umbrella type “neural tube”. We then pruned from our trees each leaf labeled with a cell type not included in our analysis (aPSM, pPSM).

**Table 1.** Summary statistics on post-processing cell lineage trees from TLS.

| # After Pre-Processing | Trees | Cell Types (Max) | Cell Types (Min) | Cell Types (Avg) | Cells (Max) | Cells (Min) | Cells (Avg) | Edges (Max) | Edges (Min) | Edges (Avg) |
| --- | --- | --- | --- | --- | --- | --- | --- | --- | --- | --- |
| TLS | 14 | 6 | 3 | 4.79 | 1723 | 15 | 336.36 | 2267 | 23 | 468.50 |

As a pre-processing step to *Carta* only, we collapsed each clade in each cell lineage tree comprised of extant cells that share a cell type into a single extant cell with that cell type. These clades do not contribute to cell type transitions nor the objective score of *Carta*.

#### 4.4.2 Processing of data in Weinreb *et. al* by study

We obtained the *in vitro* differentiation time course data generated by Weinreb *et. al* [30] from their public repository (https://github.com/AllonKleinLab/paper-data/tree/master/Lineage_tracing_on_transcriptional_landscapes_links_state_to_fate_during_differentiation). The associated metadata includes the lentiviral barcode and cell type of each cell. Each of the 5864 barcodes corresponds to a star-shaped cell lineage tree, where the leaves represent the sequenced cells that contain that barcode and are annotated by cell types (Section 2.4). Of the 130887 cells in the dataset, 49297 have an associated barcode. We observed 107 distinct potencies in the data, defined by the set of cell types of the descendants of a cell, even though the data only has 9 mature cell types. This is possibly due to cell sampling limitations, as illustrated in Section 2.1. As such, we performed a mild filtering of the data by removing barcodes in which the observed potency occurs less than 10 times in the data. This step removes only 4.1% of the barcodes, resulting in 5642 cell lineage trees totaling 43670 cells. We merged the “pDC” and “Ccr7 DC” cell types into one “DC” cell type, and removed cells with the undifferentiated cell type from the cell lineage trees. These cell lineage trees are provided as input for *Carta* and the other existing methods.

### 4.5 Implementation and application of existing methods

#### 4.5.1 Fitch’s Algorithm

Fitch’s Algorithm solves the small parsimony problem [75] which can be applied to lineage tracing data to build cell differentiation maps under the assumption that all the progenitor cell types are observed in the data. Briefly, given a phylogeny with each leaf labeled with one of a set of states, the small parsimony problem seeks to find the labeling of internal nodes of a phylogeny with those states such that the fewest number of transitions in state between parent and child nodes is obtained [75]. The frequency of transition from cell type *i* to cell type *j* can then be counted as the number of transitions from an internal cell labeled *i* to one labeled *j* in this labeled phylogeny.

For the dataset from Weinreb *et. al* (Section 2.4), we directly applied Fitch’s Algorithm and totaled the number of transitions between cell types across the Fitch labeling for each star-shaped cell lineage tree. We then stored these totals in a cell type transition matrix and row-normalized the matrix, converting transition frequencies to transition proportions that sum to 1 for each cell type of origin.

For the TLS dataset (Section 2.3), to account for the often large number of equally parsimonious Fitch labelings for large trees, we used *FitchCount* [38], which efficiently counts the total number of transitions between cell types in all equally minimal Fitch labelings. As the total number of transitions counted by FitchCount increases rapidly by the size of the cell lineage tree, the transition counts on large trees would dominate the transition count totaled over all trees. Thus, we computed a normalized sum of the transitions over all trees. For each tree we generated a row-normalized cell type transition matrix from the FitchCount transitions, and then computed the sum of these matrices as the final cell type transition matrix. This final matrix is then row-normalized.

#### 4.5.2 Evolutionary Coupling (EvoC)

Evolutionary Coupling is defined as the normalized phylogenetic distance between any pair of cell annotations on a tree [39]. We extend the definition given in Yang *et al*. [39] to multiple trees T. Given cell types *M* and *K*, the average phylogenetic distance between leaves (extant cells) labeled by these cell types on the cell lineage tree is defined as:

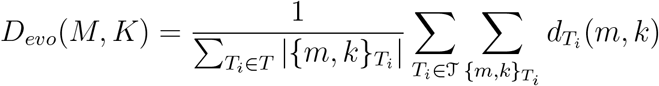

Where 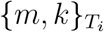 is the set of all pairwise combinations of leaves with type *M* and *K* on tree *T*_*i*_ and 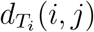 denotes the phylogenetic distance between leaves on tree *T*_*i*_. Intuitively, this metric calculates the average phylogenetic distance between two cells of cell types *M* and *K*. We then perform hierarchical clustering on the cell types based on *D*_*evo*_ using the UPGMA (Unweighted Pair Group Method with Arithmetic Mean) algorithm [80], yielding a tree-structure cell differentiation map.

#### 4.5.3 ICE-FASE

ICE-FASE calculates the average times at which cell types separate across given time-resolved cell lineage trees, and performs hierarchical clustering between these cell types to form the resultant cell differentiation map [47]. To run ICE-FASE, we used the implementation in the *QFM* package in R [47]. In addition to cell type annotations, ICE-FASE requires time-resolved phylogenies with branch lengths as input. For the TLS cell lineage trees, we estimated the branch lengths using the Maximum Likelihood Branch Length Estimator implemented in Cassiopeia [40]. The lineage tracing data from Weinreb *et al*. [30] is already annotated with time.

We implemented several workarounds in the analysis of both datasets due to limitations in the ICE-FASE codebase. Firstly, since the ICE-FASE code crashes when multiple trees are given as input, we created a single tree by connecting the root of each input tree by a 0-length branch to a dummy root node. Secondly, the ICE-FASE code is not equipped to handle trees that have polytomies, i.e. vertices with more than two children. Since the TLS trees (Section 2.3) and the Weinreb *et. al* trees (Section 2.4) both have such polytomies, we arbitrarily binarized these trees by creating edges with 0-length. Importantly, since ICE-FASE depends only on the timing at which cells separate, the introduction of these 0-length branches should not affect the analysis. Moreover, combining multiple trees into a single tree should not be problematic, as all pairs of cells in different trees now connected by the dummy root have a separation time of 0.

#### 4.5.4 PhyloVelo

PhyloVelo attempts to learn the differentiation trajectories of a system from gene expression data that is informed by the lineage depth of each cell. To run PhyloVelo, we utilized the PhyloVelo package as provided in [52]. We performed the analysis very closely to the analysis of PhyloVelo performed in that study. For both datasets, we utilized the “velocity inference” and “velocity embedding” embedding functions to calculate the PhyloVelo trajectories, and then passed the output of these functions to the “state graph” function in Dynamo [81] to obtain the cell type transition matrix. We then transposed this matrix as PhyloVelo reverses directionality in its transitions, and row normalized it as well.

For the TLS dataset (Section 2.3), we provided an anndata object generated by a standard Seurat RPCA integration pipeline of the scRNA-seq data for the sequenced TLS experiments [29]. This pipeline normalizes counts for 22291 genes and generates UMAP coordinates for each cell. We subsetted the anndata object to cells that are in the cell lineage trees, and calculated the depth of each cell as the number of edges from the root that have at least one mutation. We further removed genes with a count lower than 50 across all cells.

For the dataset from Weinreb *et. al* (Section 2.4), we generated an anndata object using the normalized gene counts from the publicly available data in the original study [30]. We included only cells with barcodes. We then closely followed the analysis suggested in the documentation of [52] (https://phylovelo.readthedocs.io/en/latest/notebook/in_vitro_hematopoiesis.html), using largely the same parameter choices. One notable difference is we used *n neigh* = 500 in the “velocity embedding” function, as using the originally specified 100 generates an error in state graph construction.

### 4.6 Choosing the optimal number of progenitors in *Carta* for real data

We selected the number *k*^*^ of progenitors by finding a elbow in the *k* vs. minimum discrepancy graph, using the *kneedle* algorithm. Initially, the *kneedle* algorithm found elbow points with very few progenitors (*k* = 4 for the DAG curve for the TLS data (Section 2.3) and *k* = 3 for the tree curve for the data from Weinreb *et. al* (Section 2.4), respectively). These elbows provided cell differentiation maps that included too few progenitors to fully capture the complex dynamics in the developmental systems we explored. We found kneedle to be conservative, selecting an elbow at the first point with a significant reduction in the difference in discrepancy with the previous point. Hence, we sought to select an elbow amongst the “flat” region of each curve to determine which progenitors whose inclusion yields the lowest value in terms of reduced discrepancy while maintaining a useful number of progenitors. Thus we applied kneedle to the regions where the curve flattens out (*k* = 4, … 11 for the DAG curve for the TLS data and *k* = 5, … 11 for the tree curve for the data from Weinreb *et. al*), giving elbows at *k* = 7 for both datasets *et. al*.

### 4.7 Choosing the edges in the cell differentiation maps inferred by *Carta* for real data

For the *Carta*-DAG cell differentiation map with *k* = 7 inferred for TLS (Figure 3d) (Section 2.3), we include all transitions that appear frequently in the cell lineage trees. Specifically, we define the *cellular flow w*(*P, P* ^*′*^) for a transition (*P, P* ^*′*^) as the number of cells across the given set of cell lineage trees that traverse through that transition. To calculate the cellular flow for a transition, we counted the instances in which ℓ(*v*) = *P*, ℓ(*u*) = *P* ^*′*^ for each edge (*u, v*) in a cell lineage tree *T*, weighting by the number of leaf descendants of *v*. This weighting preserves flow in the map such that the cellular flow entering a progenitor is equal the the cellular flow exiting it, i.e.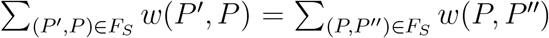. We keep an edge in the cell differentiation map in Figure 3d if: (1) the edge is necessary to ensure that an extant state is reachable by a progenitor that includes that in its potency; or (2) the edge has a cellular flow that is > 0.2\**deg*^+^(*P*) meaning that the edge accounts for more than 20% of the cellular flow from its parent progenitor. Note that this criterion also removes 0-flow edges. For the *Carta*-tree cell differentiation map with *k* = 7 inferred for the data from Weinreb *et. al* (Section 2.4), we only include edges such that the the map has a tree structure.

### 4.8 Discrepancy of differentiation maps inferred by existing methods

The first step in calculating the discrepancy of cell differentiation maps inferred by existing methods is determining the potencies of progenitors in the inferred cell differentiation maps. For methods that produce binary tree-structured cell differentiation maps (ICE-FASE and EvoC), the potency of a progenitor – i.e. an internal vertex *v* of the map – is the set of observed cell types – i.e. leaves – in the subtree rooted at *v*. As Fitch and PhyloVelo do not explicitly infer progenitors, we devise a scheme to obtain progenitors from their cell differentiation maps. The output of Fitch and PhyloVelo is a normalized transition frequency (*f* (*t*_*i*_, *t*_*j*_)) between each pair of states *t*_*i*_, *t*_*j*_ ∈ *S*. For each observed cell type *t*_*i*_, we introduce a progenitor as {*t*_*j*_ : *f* (*t*_*i*_, *t*_*j*_) ≥ *ϵ*}. This is the set of each cell type *j* for which the transition frequency from a cell type *i* exceeds threshold *ϵ*. In this work we chose 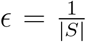, and thus *ϵ* = 0.166 for the TLS data (Section 2.3) and *ϵ* = 0.111 for the data from Weinreb *et. Al* (Section 2.4).

We computed the discrepancy for each method by solving the Progenitor Labeling Problem (PLP; Problem A.1) using the dynamic programming algorithm outlined in Supplementary Section A.3. The number of cell type transitions (Figure 3f,g) are determined by the inferred minimum discrepancy labeling of the cell lineage trees.

### 4.9 Calculating distances for undifferentiated cells in the data from Weinreb *et. al*

We labeled each undifferentiated cell (cells labeled with the “undifferentiated” cell type in [30]) in the data from Weinreb *et. al* (Section 2.4) with the progenitor type that *Carta* assigns to the ancestral cell of its star-shaped cell lineage tree – these are the labels shown in Figure 5a. We next describe how we calculate the distance of each undifferentiated cell to the closest observed cell type cluster in gene-expression space. First, we obtained the normalized counts for 25289 genes across all cells in this dataset from the publicly available *in vitro* differentiation time course data from Weinreb *et. al* [30] (https://github.com/AllonKleinLab/paper-data/tree/master/Lineage_tracing_on_transcriptional_landscapes_links_state_to_fate_during_differentiation). We then removed cells with counts = 0 or counts >1, 000, 000 and performed principal components analysis with *n* = 50 components. We next calculated a 50 PC centroid for each observed cell type by averaging across the PC values of cells of that type, and then calculated the euclidean distance in PC values between each undifferentiated cell and each centroid. Finally, in Figure 5c, for each observed cell type cluster, we calculated the proportion of undifferentiated cells labeled with each progenitor cell type by *Carta* that is closest to that cluster.

## Acknowledgements

This work is supported by grant 2024-345885 from the Chan Zuckerberg Initiative DAF, an advised fund of Silicon Valley Community Foundation; NIH grant DP2HD111537 to M.M.C.; NCI grant U24CA248453 to B.J.R; and the Princeton Catalysis Initiative. R.Z. is supported by T32HG003284.

## Competing Interests

The authors declare no competing interests.

## Author Contributions

P.S., R.Z., M.M.C. and B.J.R. conceived and developed the method. P.S. and R.Z. implemented the software and performed analysis of the lineage tracing data. B.L. produced the cell lineage trees and cell type annotation for the TLS data. A.S. ran competing methods on real data. P.S. and H.S. derived the complexity proofs. P.S., R.Z., A.B., M.M.C. and B.J.R. interpreted the results and wrote the manuscript. All authors read and approved the manuscript.

## Data Availability

Simulated data is available on github at https://github.com/raphael-group/CARTA/tree/main/data. Raw sequencing data for the trunk-like-structures dataset is available at https://www.ncbi.nlm.nih.gov/geo/query/acc.cgi?acc=GSE220949. The processed lineage trees and cell type annotations are available at https://github.com/raphael-group/CARTA/tree/main/data/gastruloid. The single-cell lineage tracing data of mouse hematopoiesis is available at https://github.com/AllonKleinLab/paper-data/tree/master/Lineage_tracing_on_transcriptional_landscapes_links_state_to_fate_during_differentiation.

## Code Availability

The code is publicly available at https://github.com/raphael-group/CARTA under BSD 3-Clause license.

## A Supplementary Information

### A.1 Definition of discrepancy

Let 𝒯 := {*T*_1_, …, *T*_*m*_} be *m* cell lineage trees, each describing the cell division history of a distinct biological replicate that belong to the same developmental system. Let *F*_*S*_ be a cell differentiation map for the set *S* of observed cell types. *F*_*S*_ is a directed graph whose vertices represent observed cell types and unobserved progenitors – and are labeled by either an elements of *S or* a subset of *S* – and whose edges represent cell type transitions that occurred during development.

We first define a progenitor labeling 𝓁 that, for a given cell differentiation map *F*_*S*_, labels the cells (vertices of 𝒯) with cell types (vertices of *F*_*S*_). Let 𝓁_*t*_ denote the cell type labeling of the leaves of cell lineage trees 𝒯. The progenitor labeling must follow constraints imposed by the differentiation map and cell lineage trees. First, the progenitor type of a cell must contain the cell types of the descendants of the cell in the cell lineage tree. More formally, we require that for each vertex *v* of tree *T*∈ 𝒯, 𝓁(*v*) ⊇*B*(*v*), where *B*(*v*) is the set of observed cell types of the leaves in the subtree of *T* rooted at vertex *v*. Second, since the cells differentiate under the model described by the cell differentiation map *F*_*S*_, for any edge (*u, v*) ∈*E*_*T*_ in a lineage tree *T* ∈𝒯, there must be a directed path from the progenitor 𝓁(*u*) to the progenitor 𝓁(*v*) in the cell differentiation map *F*_*S*_. We formally describe these conditions on the progenitor labeling as follows.

#### Definition A.1.

*A progenitor labeling 𝓁 is* compatible *with a cell lineage tree 𝒯 and a cell differentiation map F*_*S*_ = (𝒫_*S*_, *E*_*F*_) *if and only if (i) 𝓁*(*v*) ∈ 𝒫 *for each vertex v of T; (ii) 𝓁*(*v*) ⊇ *B*(*v*) *for each vertex v of T; (iii) there is a directed path from 𝓁*(*u*) *to 𝓁*(*v*) *in F*_*S*_ *for each edge* (*u, v*) *in T*.

For cell lineage trees 𝒯 and a cell differentiation map *F*_*S*_, there may be multiple compatible progenitor labelings 𝓁. Each compatible labeling 𝓁 induces a discrepancy *D*(𝒯, *F*_*S*_, 𝓁) between the trees 𝒯 and the map *F*_*S*_ which arises from two limitations of current lineage tracing data.

First, all of the observed cell types in the potency of each cell will not necessarily be observed in each cell lineage. This may lead to a mismatch between the potency 𝓁(*v*) of an ancestor cell *v* and the observed cell types *B*(*v*) of its descendants. We quantify this mismatch by the sampling discrepancy *D*_*s*_(𝒯, *F*_*S*_, 𝓁) which penalizes the absence of feasible cell types in the descendants of a progenitor as,

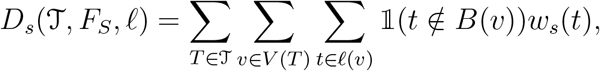

where *w*_*s*_(*t*) is the weight associated with cell type *t* that can reflect the relative sampling abundance of different cell types. In this work, we use *w*_*s*_(*t*) = 1 for all *t*.

Second, instances in which cell divisions that are accompanied by a differentiation event are not resolved in the inferred cell lineage tree will lead to discrepancy *D*_*r*_(𝒯, *F*_*S*_, 𝓁) between the observed and the true transitions between cell types during development. We call this *resolution discrepancy* and quantify it as,

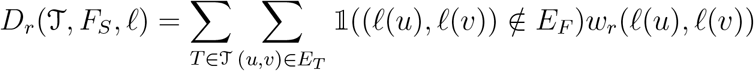

where *w*_*r*_(*P, P* ^*′*^) is the weight associated with cell type transition (*P, P* ^*′*^) that can reflect the likelihood of that transition occurring. We introduce a parameter *λ* to reflect the relative likelihood of the occurrence of sampling and resolution discrepancies, and define the total discrepancy *D*(*T, F*_*S*_, 𝓁) as follows.

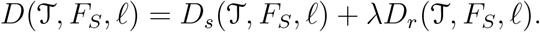

We refer to the problem of finding a progenitor labeling 𝓁 that induces the minimum discrepancy between cell lineage trees 𝒯 and cell differentiation map *F*_*S*_ as the Progenitor Labeling Problem (PLP), which we define as follows.

#### Problem A.1

(Progenitor Labeling Problem (PLP)). *Given a set* 𝒯 *of cell lineage trees and cell differentiation map F*_*S*_, *find a progenitor labeling 𝓁 that is compatible with* 𝒯 *and F*_*S*_, *and minimizes the discrepancy D*(𝒯, *F*_*S*_, 𝓁).

This is an analog of the *small parsimony* problem [75], and we show that this problem can be solved by a dynamic program by adapting Sankoff’s algorithm [76] (Supplementary Section A.3).

We define the discrepancy *D*(𝒯, *F*_*S*_) between cell lineage trees 𝒯 and a cell differentiation map *F*_*S*_ as

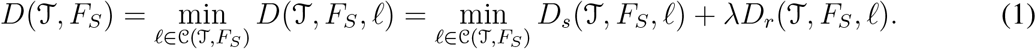

where 𝒞 (𝒯, *F*_*S*_) is the set of progenitor labelings that are compatible with cell lineage tree *T* and cell differentiation map *F*_*S*_. As discussed, naively minimizing the discrepancy may lead to inference of a large cell differentiation map with false positive progenitors and cell type transitions (Figure 1c). To prevent this, we impose constraints on the size of the cell differentiation map. Specifically, we find a cell differentiation map with *k* progenitors, and *k*^*′*^ edges (cell type transitions) that has the minimum discrepancy with given cell lineage trees 𝒯.

The discrepancy *D*(𝒯, *F*_*S*_) between cell lineage trees 𝒯 and a cell differentiation map *F*_*S*_ depends on the choice of *λ* (Equation 1), which reflects the relative likelihood of the sampling and the resolution discrepancies. Most current lineage tracing technologies, such as CRISPR-Cas9 based lineage tracers [28, 39, 40], utilize a limited number (10-30) of sites where mutations are induced and each site can acquire only one mutation along a lineage in the cell lineage tree. This limits the depth and resolution of the reconstructed cell lineage tree, leading to several resolution discrepancies. In contrast, a sampling discrepancy of a cell type in the potency of a progenitor cell only occurs when none of the descendants with that cell type are sequenced. Given the high throughput (thousands of cells) [28, 30, 36] of current lineage tracing technologies, this makes the occurrence of resolution discrepancies much more likely compared to sampling discrepancies. As such, in this study, we set the parameter *λ* = 0 to reflect the relative likelihood of the two kinds of discrepancies.

### A.2 *Carta*: mixed integer linear programming (MILP)

#### A.2.1 *Carta*-DAG MILP formulation

To solve the CDMIP problem, for a given cell lineage trees 𝒯 the *Carta*-DAG MILP finds a cell fate map *F*_*S*_ with *k* progenitor cell types and the progenitor labeling 𝓁 with the minimum induced discrepancy *D*_*s*_(𝒯, *F*_*S*_, 𝓁).

##### Progenitor labeling

We start by introducing binary variables *x*_*p,t*_ for *p* ∈ {1, …, *k*} and *t* ∈ *S* to represent the set of *k* progenitors in the cell fate map, such that *x*_*p,t*_ = 1 if cell type *t* is in the potency of progenitor *p* and *x*_*p,t*_ = 0 otherwise. The assignment of each vertex to one of the progenitors is encoded by binary variables *y*_*v,p*_ for each vertex *v* in cell lineage tree *T* and progenitor *p*, such that *y*_*v,p*_ = 1 if *v* is labeled by *p*. Since each vertex *v* can only be labeled by one progenitor, we enforce the following constraint for each vertex *v*.

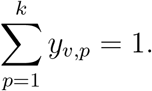

##### Compatibility constraints

We must enforce constraints to ensure compatibility of the progenitor labeling with the cell lineage tree and the cell fate map (Definition A.1). We require that (i) if vertex *v* is labeled by progenitor *p* then *p* must have the potency for all the cell types in *B*(*v*) and (ii) for each edge (*u, v*) in each tree *T* ∈ 𝒯, progenitor 𝓁(*v*) is reachable from progenitor 𝓁(*v*) in *F*_*S*_.

We introduce continuous variables *z*_*v,t*_∈ [0, 1] for each vertex *v* of each tree *T* ∈𝒯 and cell type *t* to indicate if cell type *t* belongs in the potency of progenitor 𝓁(*v*). Specifically, we introduce constraints to ensure that *z*_*v,t*_ = 1 if *t* ∈ 𝓁(*v*) and *z*_*v,t*_ = 0 otherwise. We achieve this by enforcing,

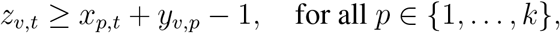

each vertex *v* and cell type *t* to make sure that *z*_*v,t*_ = 1 if *x*_*p,t*_ = 1 and *y*_*v,p*_ = 1 for some progenitor *p*. Note that we do not need to introduce additional constraints to bound *z*_*v,t*_ from above since minimizing the sampling discrepancy will ensure that *z*_*v,t*_ = 0 if either *x*_*p,t*_ = 0 or *y*_*v,p*_ = 0 for all *p* ∈ {1, …, *k*}.

We ensure that the progenitor label of each vertex *v* contains *B*(*v*) by encoding the following constraints,

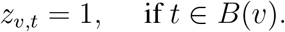

Further, for each edge (*u, v*) of *T* ∈𝒯, we introduce the following constraints that ensure that the progenitor label of *u* contains the progenitor label of *v*,

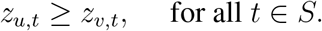

##### Objective function

Our goal is to find a cell fate map that minimizes *D*(𝒯, *F*_*S*_) for cell lineage trees 𝒯. This is equivalent to finding a cell fate map *F* _*S*_ and progenitor labeling 𝓁 such that the induced discrepancy *D*_*s*_(𝒯, *F*_*S*_, 𝓁) is minimized. In the case of *λ* = 0, we only need to minimize the induced sampling discrepancy which can be described in terms of variables *z*_*v,t*_ as,

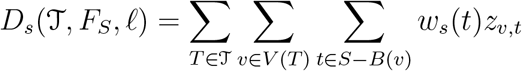

where *w*_*s*_(*t*) is the penalty of not sampling cell type *t*. Note in this work we use *w*_*s*_(*t*) = 1 for all *t*. We minimize the above objective in the MILP that has *O*(*n* |*S* |+ |*S*| *k*) binary variables, *O*(*n*| *S* |) continuous variables and *O*(*n*| *S* |*k*) constraints, where *n* is the total number of vertices across all cell lineage trees.

#### A.2.2 *Carta*-tree MILP formulation

To solve the CDTIP problem (Problem 4.3), we show that there is an equivalence (Theorem A.1) between the CDTIP problem (Problem 4.3) and the Weighted Insertion Flip Problem (WIFP) (Problem A.2) (see Supplementary Section A.4). We thus formulate an mixed integer linear problem (MILP) for the WIFP problem to solve the CDTIP problem.

We introduce binary variables 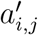 for each row *i* and column *j* to encode the binary matrix *A*^*′*^. Since we only allow 0 → 1 flips, we enforce that 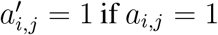 using the following constraint,

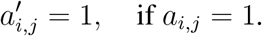

We require that *A*^*′*^ must admit a perfect phylogeny. We use the set inclusion and disjointness (SID) formulation described in Chimani *et al*. [82]. This formulation uses a characterization of perfect phylogeny matrices that states that for any two columns, we require the 1-sets, i.e. set of rows *i* such that 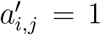, of any two columns should either be disjoint or related by containment [83] (Theorem A.3). We introduce continuous variables *y*_*j,j*_*′* and *z*_*j,j*_*′* for each pair of columns *j* and *j*^*′*^. We force *y*_*j,j*_*′* = 0 if the 1-set of column *j* is not contained in 1-set of column *j*^*′*^ using the following constraint for each row *i*,

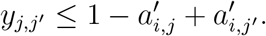

Along the same vein, we enforce *z*_*j,j*_*′* = 0 if the 1-set of column *j* is not disjoint with 1-set of column *j*^*′*^ using the following constraint for each row *i*,

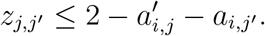

Finally, we enforce that for any two columns *j* and *j*^*′*^, at least one of *y*_*j,j*_*′, y*_*j,j*_ *′* and *z*_*j,j*_*′* must be 1 using the following constraint.

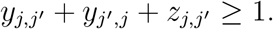

We penalize the weighted sum of the 0 → 1 flips by minimizing the following objective function,

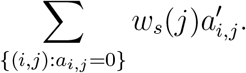

### A.3 Dynamic programming algorithm to solve the PLP problem

We consider a single tree *T*∈ 𝒯. For differentiation map *F*_*S*_, let *D*_*s*_(*T* (*v*), *F, 𝓁*(*v*)) be the number of discrepancies in the subclade of *T* rooted at vertex *v* if it has progenitor label 𝓁(*v*) ∈ *F*_*S*_. For each *v* that is a leaf of *T*, we initialize *D*_*s*_(*T* (*v*), *F*_*S*_, 𝓁(*v*)) as 0 if the observed state at *v* is 𝓁(*v*) and ∞ otherwise.

The recurrence relation is:

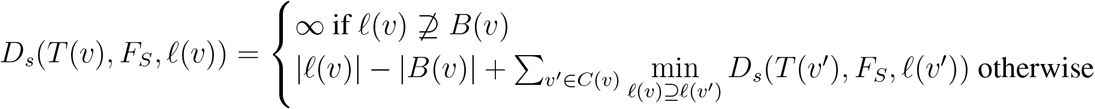

Where *C*(*v*) denotes the direct descendants of *v* in *T*.

We can then find the total discrepancy over all cell lineage trees as 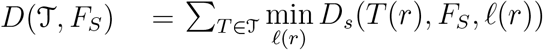, *F*_*S*_, 𝓁(*r*)) where *r* is the root of *T*.

The labeling 𝓁 that minimizes *D*_*s*_(*T, F*_*S*_, 𝓁(*v*)) for each tree *T* can then be acquired by taking a subsequent top-down pass through *T* as in Sankoff’s algorithm. Start by labeling *r* as 𝓁(*r*) = min *D*_*s*_(*T* (*r*), *F*_*S*_, 𝓁(*r*)). Then for each vertex *v*^*′*^, let 𝓁(*v*) be the chosen progenitor label of parent *v* in *T*, and take 𝓁(*v*^*′*^) as 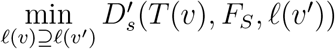.

This algorithm operates in *O*(*nm*) time, where *n* is the number of vertices across all cell lineage trees in 𝒯 and *m* is the number of vertices in *F*_*S*_.

### A.4 Equivalence of Cell Differentiation Tree Inference Problem and the Weighted Insertion Flip Problem

To derive a characterization of cell differentiation trees (cell differentiation maps constrained to have tree structures), we draw a connection to two-state perfect phylogenies [84]. Specifically, we show that the Cell Differentiation Tree Inference Problem (Problem 4.3) is equivalent to a weighted version of the Minimum Insertion Flip Problem [85].

We begin by showing that when the cell differentiation map is a tree, the Progenitor Labeling Problem (PLP) (Problem A.1) can be solved more efficiently than in the general case when the cell differentiation map is a directed acyclic graph (DAG). The Progenitor Labeling Problem (PLP) is a special case of the small parsimony problem in phylogenetics [75] and can be solved using dynamic programming by adapting the Sankoff’s Algorithm [76] (details in Section A.3). However, when the cell differentiation map is a tree, the PLP problem can be solved even more efficiently by labeling each vertex *v* of the cell lineage tree *T* by the smallest feasible progenitor. This labeling 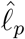is obtained if we label each vertex *v* by the lowest common ancestor (LCA) of the cell types in the observed potency *B*(*v*) in the cell differentiation tree *F*_*S*_. More formally, we have the following lemma.

#### Lemma A.1.

*Given lineage trees* 𝒯 *and a cell fate tree F*_*S*_, *the progenitor labeling* 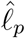 *that labels each vertex v by the lowest common ancestor (LCA) of the observed potency B*(*v*) *in F*_*S*_ *yields the minimum induced discrepancy, i*.*e*. 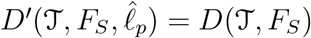.

The proof for this lemma is given in Supplementary Section A.6.1.

Next, we derive the constraints on the set of progenitors in a cell differentiation tree. Specifically, we show that the set of progenitors in a cell differentiation tree must be *laminar*, i.e. the potencies of each pair of progenitors are either disjoint or related by containment.

#### Lemma A.2.

*The set* 𝒫 *of progenitors in a cell differentiation tree F*_*S*_ *is laminar, i*.*e. for any two distinct progenitors 𝒫 and P* ^*′*^, *either P* ∩ *P* ^*′*^ = ∅ *or P* ⊂ *P* ^*′*^ *or P* ^*′*^ ⊂ *P*.

The proof for this lemma is given in Supplementary Section A.6.2.

Laminar set families are closely connected to perfect phylogenies [84]. A phylogeny is a tree whose leaves represent the extant taxa and the internal vertices represent ancestral taxa, and is called a *perfect phylogeny* if each character changes states according to the infinite sites model [86]. Specifically, each character is allowed to change from state 0 to state 1 exactly once and never changes from state 1 to state 0 along the edges of the phylogeny. The taxa can be represented by a binary matrix, called the *character matrix*, where each row is a taxon and each column is a character. It has been shown that a character matrix admits a perfect phylogeny if and only if the sets of taxa with state 1 for each character form a laminar family of sets [84]. As such, not all character matrices will admit a perfect phylogeny and a standard problem in phylogenetics, called the *Minimum Flip Problem* [85], is to find the smallest number of flips (both 0 →1 and 1 →0) that must be made in a given character matrix for it to admit a perfect phylogeny. When only 0 →1 flips are allowed, this problem is referred to as the *Insertion Flip Problem* (IFP) [85, 87]. We formally pose the weighted version of the IFP problem as follows.

#### Problem A.2

(Weighted Insertion Flip Problem (WIFP)). *Given a m × n binary matrix A, find binary matrix A*^*′*^ *by only doing* 0 → 1 *flips such that the following weighted sum of the flips*

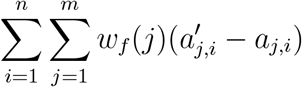

*is minimized, where w*_*f*_ (*j*) *is the weight associated with flipping an entry in row j*.

The IFP problem is a special case of the WIFP problem where *w*_*f*_ (*j*) = 1 for all rows *j* ∈ {1, …, *m*}.

We derive a connection between the CDTIP problem (Problem 4.3) and the Weighted Insertion Flip Problem (WIFP) using Lemma A.1 and Lemma A.2. Specifically, we show that any instance of the CDTIP problem can be transformed in polynomial time to an instance of the WIFP problem.

#### Theorem A.1.

*The CDTIP and WIFP problems are m-equivalent, i*.*e. these problems are polynomial time mapping reducible to each other*.

The proof for this theorem is given in Supplementary Section A.6.3.

Theorem A.1 implies that algorithms and heuristics that have been developed to solve the WIFP problem [87, 88] can be used to solve the CDTIP problem.

### A.5 Characterizing the complexity of the CDTIP and CDMIP problems

Since it has been shown that the IFP problem is NP-complete [85], as a corollary of Theorem A.1, we have that CDTIP problem is also NP-complete.

#### Corollary A.1.

*The CDTIP problem is NP-complete*.

Interestingly, since the CDTIP problem minimizes the discrepancy between multiple trees, namely the cell lineage trees 𝒯 and the cell differentiation tree *F*_*S*_, it belongs to the large class of problems called the *Tree Reconciliation Problem* [89]. In the Tree Reconciliation Problem, the discrepancy between trees is quantified using a mapping between the vertex sets of the trees, which in our case is the progenitor labeling 𝓁.

Using a parsimonious reduction from the Minimum Vertex Cover problem [90], we also show that the more general CDMIP problem, where the cell differentiation map is a directed acyclic graph, is also NP-hard and counting the number of solutions of the CDMIP problem, which we refer to as the #CDMIP problem, is #P-hard.

#### Theorem A.2.

*The CDMIP problem is NP-hard and the #CDMIP problem is #P-hard*.

The proof for this theorem is given in Supplementary Section A.6.4.

### A.6 Proofs

#### A.6.1 Proof for Lemma A.1

Here, we provide a proof for Lemma A.1. Specifically, we show that for a given cell lineage tree *T* and cell fate tree *F*_*S*_, the progenitor labeling 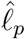 that labels each vertex *v* by the lowest common ancestor (LCA) of the observed potency *B*(*v*) in *F*_*S*_ yields the minimum induced discrepancy, i.e. 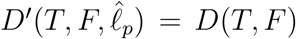. Recall that observed potency *B*(*v*) for a vertex *v* of lineage tree *T* is defined by the set of cell types of the leaves in the subtree of *T* rooted at *v*. We re-state Lemma A.1 here for completeness and then provide the proof.

##### Lemma A.1.

*Given lineage trees* 𝒯 *and a cell fate tree F*_*S*_, *the progenitor labeling* 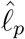 *that labels each vertex v by the lowest common ancestor (LCA) of the observed potency B*(*v*) *in F*_*S*_ *yields the minimum induced discrepancy, i*.*e*. 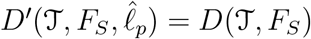.

*Proof*. Since each vertex is labeled by the smallest feasible set of cell types that contains the observed potency, labeling 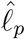 clearly minimizes the sampling discrepancy which penalizes the deviation of the progenitor from the observed potency. To complete the proof, we must show that progenitor labeling 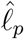 is compatible with the cell lineage tree *T* and cell fate tree *F*_*S*_ (Definition A.1). There are three conditions of compatibility. By definition of 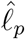, it is clear the 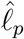 satisfies condition (i), which states that 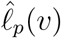 must be a progenitor in the cell fate tree *F*_*S*_ for each vertex *v*, and condition (ii), which states that 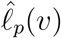 must contain *B*(*v*) for each vertex *v*. We now show that 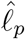 satisfies condition (iii), which states that 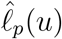 must contain 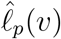 for each edge (*u, v*) in *T*. Since (*u, v*) is an edge of *T*, we have *B*(*u*) ⊃ *B*(*v*). Since *F*_*S*_ is a tree, this implies that lowest common ancestor of *B*(*u*) precedes the lowest common ancestor of *B*(*v*). Therefore, 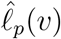 is reachable from 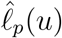 in *F*_*S*_. □

#### A.6.2 Proof for Lemma A.2

Here we show that the progenitors in a cell differentiation tree are laminar, i.e. the potencies for each pair of progenitors are either disjoint or related (Lemma A.2). For completeness, we restate the lemma here and then provide a proof.

##### Lemma A.2.

*The set* 𝒫 *of progenitors in a cell differentiation tree F*_*S*_ *is laminar, i*.*e. for any two distinct progenitors 𝒫 and P* ^*′*^, *either P* ∩ *P* ^*′*^ = ∅ *or P* ⊂ *P* ^*′*^ *or P* ^*′*^ ⊂ *P*.

*Proof*. We prove this by contradiction. Suppose there are two progenitors *P* and *P* ^*′*^ that are neither disjoint nor related by containment. Then, there exists distinct cell types *a, b* and *c* such that *a* ∈*P \P* ^*′*^, *b* ∈*P \P* ^*′*^ and *c* ∈*P* ^*′*^*\ P*. By definition of cell differentiation map there must exist a path from progenitor *S* (totipotent cell) to {*b*} going through *P* and a path from *S* to {*b*} going through *P* ^*′*^. Since *P* and *P* ^*′*^ are not related by containment, this violates the premise that *F*_*S*_ is a tree.

#### A.6.3 Proof for Theorem A.1

As we have seen in Lemma A.1 that the labeling 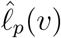 that induces the minimum discrepancy between a cell fate tree *F*_*S*_ cell lineage tree *T* completely characterized by the observed potency *B*(*v*). We use this property to show that the WIFP and the CDTIP problems are equivalent.

We start by stating the following well-known characterization of perfect phylogeny matrices [83].

##### Theorem A.3.

*Suppose A* ∈ {0, 1}^*n×m*^ *is a binary character-matrix and I*(*j*) := {*i* : *a*_*i,j*_ = 1, *i* ∈ [*n*]*} is the set of indices of the ‘1’ entries in the j*^*th*^ *column of A. A admits a perfect phylogeny if and only if for each pair j, j*^*′*^ *of characters, either: (i) I*(*j*) ⊆ *I*(*j*^*′*^); *(ii) I*(*j*^*′*^) ⊆ *I*(*j*); *or (iii) I*(*j*) ∩ *I*(*j*^*′*^) = ∅.

This theorem states that the a binary matrix *A* is a perfect phylogeny matrix if and only if the one-sets *I*(*j*) of the columns *j* ∈ {1, …, *m*} of *A* are laminar.

We restate the theorem for completeness and then proceed with a proof.

##### Theorem A.1.

*The CDTIP and WIFP problems are m-equivalent, i*.*e. these problems are polynomial time mapping reducible to each other*.

*Proof*. We start by showing a polynomial-time reduction from the WIFP problem to the CDTIP problem.

Let *m× n* binary matrix *A* and weight function *w*_*f*_ form an instance of the WIFP problem. We build a corresponding instance of the CDTIP problem as follows. We start with constructing a cell lineage tree *T*. *T* has a root *r* with *n* children *v*^(1)^, …, *v*^(*n*)^, each corresponding to a row of *A*. Vertex *v*^(*i*)^ has one child *v*^(*i,j*)^ for each column *j* in *A* such that *a*_*j,i*_ = 1. As such, the tree *T* has one leaf for each entry (*j, i*) of *A* such that *a*_*j,i*_ = 1. The set *S* of observed cell types is given by {1, …, *m*} where *m* is the number of columns of *A*. We construct a weight function *w*_*s*_ for the sampling discrepancy such that *w*_*s*_(*j*) = *w*_*f*_ (*j*) for all *j*∈ *S*. We set the number *k* of progenitors as *m*. Clearly this construction can be performed in polynomial-time.

We now show that the constructed instance of the CDTIP problem admits a solution with discrepancy at most *γ* if and only if there exists a solution to the WIFP problem with total weight of flip of at most *γ*.

(⇒) Let 𝓁 be the progenitor labeling in the solution of the CDTIP problem. We get the solution *A*^*′*^ to the WIFP problem by setting 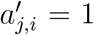 if *j* ∈ 𝓁(*v*^(*i*)^). Clearly, 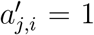 if *a*_*j,i*_ = 1, because if *a*_*j,i*_ = 1, we have *j* ∈*B*(*v*^(*i*)^) and as such *j* ∈𝓁(*v*^(*i*)^). Therefore, *A*^*′*^ is derived from *A* using only 0 → 1. Since *F*_*S*_ is a cell differentiation tree, the set of progenitors induced by 𝓁 are laminar (by Lemma A.2). Since the progenitors induced by the labeling 𝓁 correspond to one-set of columns of *A*^*′*^, by Theorem A.3 we have that *A*^*′*^ is a perfect phylogeny. Lastly, since each flip corresponds to a sampling discrepancy and *w*_*s*_(*j*) = *w*_*f*_ (*j*), the total cost of flips is equal to the total discrepancy which bounded from above by *γ*.

(⇐) Let *A*^*′*^ be the solution to the WIFP problem. We get a corresponding solution to the corresponding CDTIP problem by the progenitors induced by the labeling 𝓁, where 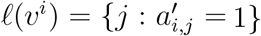. Since *A*^*′*^ is obtained from only 0 → 1 flips from *A*, we have 𝓁(*v*^(*i*)^) ⊇ *B*(*v*^(*i*)^). Since the progenitors are given by the one-sets of columns of *A*^*′*^ and *A*^*′*^ is a perfect phylogeny matrix, by Theorem A.3, we have that the progenitors are laminar. As such, the induced cell differentiation map *F*_*S*_ is a tree. Finally, we observe that the sampling discrepancy is equal to the total weight of all the flips which is *γ*. This concludes the proof.

We now show a polynomial-time reduction from the CDTIP problem to the WIFP problem. Let cell lineage tree *T* with *n* vertices, set of observed cell types *S* and integer *k* be an instance of the CDTIP problem. We construct a *m* × *n* character matrix *A* for an instance of the WIFP problem as follows. We set *a*_*j,i*_ = 1 if cell type *j* is observed in the descendants of cell *i*. As such, the one-set of column *i* is the same as the observed potency *B*(*i*) in the cell lineage tree. Additionally, we set *w*_*f*_ (*j*) = *w*_*s*_(*j*) for each cell type *j*. We show that there exists a solution *A*^*′*^ of the WIFP problem with weighted sum *γ* of the 0 → 1 flips if and only if the CDTIP problem has a solution with discrepancy *γ*.

(⇒) Let *F*_*S*_ be the solution of the CDTIP problem. Let 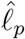 be the progenitor labeling that admits the minimum discrepancy where, from Lemma A.1, each cell *i* is labeled by the LCA of the cell types *B*(*i*) in *F*_*S*_. We construct a solution *A*^*′*^ for the WIFP problem as follows. For each column *j*, we get 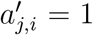 if 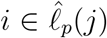. Clearly, 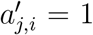 if *a*_*j,i*_ = 1 since 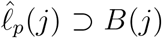. Therefore, *A*^*′*^ can be obtained from *A* by only dong 0→ 1 flips such that the total weight of the flips is *γ*. Moreover, the one-set of each column of *A*^*′*^ is one of the progenitors in *F*_*S*_ Since *F*_*S*_ is a cell differentiation tree, from Lemma A.2 the progenitors are laminar. As such, the one-set of columns of *A*^*′*^ are also laminar and therefore *A*^*′*^ is a perfect phylogeny matrix from Theorem A.3.

(⇐) Let *A*^*′*^ be the solution of the WIFP problem, i.e. *A*^*′*^ is a perfect phylogeny obtained from *A* by performing only 0→ 1 flips such that the total weight of all the flips is *γ*. Since *A*^*′*^ is a perfect phylogeny, the one-set of the columns of *A*^*′*^ are laminar and can be used to construct a cell differentiation tree *F*_*S*_. We will show that *F*_*S*_ admits a progenitor labeling such that the induced discrepancy score is *γ*. Consider the progenitor labeling 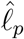 defined in Lemma A.1 where each cell of *T* is labeled by the LCA of the observed potency of the cell in *F*_*S*_. Since the one-set of *A* are the same as the observed potency of the cells in the cell lineage tree *T*, the total discrepancy induced by this labeling is *γ*. This concludes the proof.

#### A.6.4 Proof of Theorem A.2

##### Theorem A.2.

*The CDMIP problem is NP-hard and the #CDMIP problem is #P-hard*.

*Proof*. We prove the hardness of the CDMIP problem by providing a parsimonious reduction from the vertex cover problem [91]. In particular, given a *G* = (*V, E*) and a positive integer *k* ∈ N, the vertex cover problem asks if there exists a cover *C* ⊆ *V* of size at most *k* such that each edge *e* ∈ *E* has at least one endpoint in *C*.

Let *G* = (*V* (*G*), *E*(*G*)) and *k* ∈ ℕ be an instance of the vertex cover problem. We assume that there are strictly more than *k* vertices and *k* edges in *G*, since otherwise, there is a trivial vertex cover of *G*. We construct a lineage tree *T* with the following vertex set *V* (*T*), edge set *E*(*T*), and leaf labeling 𝓁_*t*_:

- A root vertex *r* ∈ *V* (*T*).
- |*E*(*G*)||*V* (*G*)| children of *r* named *v*^(*i*)^ ∈ *V* (*T*) where *v* ∈ *V* (*G*) and *i* ∈ {1, …, |*E*(*G*)|}.
- |*E*(*G*)| children of *r* named *e* ∈ *E*(*T*) where *e* ∈ *E*(*G*).
- |*V* (*G*)| − 1 children of each *v*^(*i*)^ ∈ *V* (*T*) named 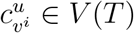 where *u* ≠*v* and 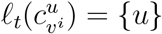.
- |*V* (*G*)| − 2 children of each *e* ∈ *V* (*T*) named 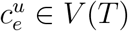 where *u* ∉ *e* and 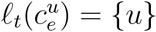.

The constructed tree *T* has depth 3, |*E*(*G*) ||*V* (*G*) | + |*E*(*G*) |+ 1 internal vertices, |*E*(*G*) ||*V* (*G*) | (*V* | (*G*) −1) + |*E*(*G*) | (*V*| (*G*) | −2) leaves, and cell type set *S* = *V* (*G*). Now, we will prove that there exists a vertex cover *C* of *G* of size at most *k* if and only if there exists a labeling 𝓁 : *V* (*T*) → 𝒫^*S*^ of *T* with discrepancy at most (|*V* (*G*)| − *k* + 1)|*E*(*G*)| such that *i) 𝓁*(*u*) ⊇ 𝓁(*v*) for all (*u, v*) ∈ *E*(*T*) and *ii)* there are at most *k* + 1 distinct labels 𝓁(*v*).

(⇒) Suppose *G* has a vertex cover *C* of size *k*. Define the labeling 𝓁 as

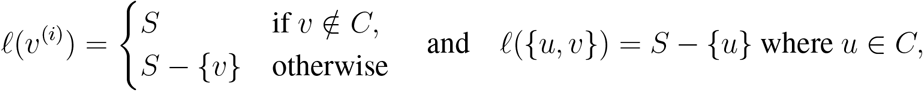

where 𝓁(*r*) = *S*. This is a valid labeling since it satisfies property *i)* and *C* is a vertex cover, which means 𝓁(*u, v*) is well-defined. Further, it uses at most *k* + 1 distinct labels as the size of *C* is less than or equal to *k*. The discrepancy of 𝓁 at the root is 0, the discrepancy of 𝓁 at each vertex *e* is 1, the discrepancy of 𝓁 at each vertex *v*^(*i*)^ where *v* ∈ *C* is 0, and the discrepancy of 𝓁 is 1 at every other vertex *v*^(*i*)^. Thus, the total discrepancy is |*E*(*G*) | + (|*V* (*G*) |−*k*) −*E*(*G*), proving the first direction of the theorem.

(⇐) Let 𝓁 : *V* (*T*) → 𝒫^*S*^ be a labeling satisfying *i)* and *ii)* with discrepancy at most (|*V* (*G*)| − *k* + 1)|*E*(*G*)|. Define the set *C* as

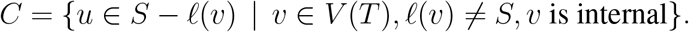

We will show that *C* is a vertex cover for *G* of size at most *k*. First, suppose that 𝓁(*u, v*) = *S* −{*u, v{* for some vertex {*u, v*}∈ *V* (*T*). Then, since there are at most *k* + 1 labels of 𝓁 and 𝓁(*r*) = *S*, there are at most *k* − 1 labels of the form *S* −{*u*}. However, this implies that at least (|*V* (*G*) | (*k* 1)) |*E*(*G*) | vertices *v*^(*i*)^ are labeled by *S*, contradicting the fact that 𝓁 has the stated discrepancy. Thus, 𝓁(*v*) has the form *S* or *S* −{*u*} for all internal vertices *v*∈ *V* (*T*). This implies that the size of *C* is at most *k*.

Now, we will show that *C* is a vertex cover of *G*. To see this, observe that the discrepancy of all vertices *v*^(*i*)^ is at least (|*V* (*G*)|−*k*)|*E*(*G*)|, since there are only *k* distinct labels of the form *S*−{*u*}. By the stated discrepancy bound, the total discrepancy of the vertices *e* can be at most |*E*(*G*)|. However, since the discrepancy of each vertex *e* is at least 1, this implies that 𝓁(*e*) = *S* − {*u*} for some *u* ∈ *e*, proving that *C* is a vertex cover of *G*.

### A.7 Additional Figures

## Notes

https://github.com/raphael-group/CARTA

### Competing Interest Statement

The authors have declared no competing interest.

## References

[1] Charles B Kimmel, Rachel M Warga, and Thomas F Schilling. Origin and organization of the zebrafish fate map. Development, 108(4):581–594, 1990.

[2] Daniel Axelrod. Carbocyanine dye orientation in red cell membrane studied by microscopic fluorescence polarization. Biophysical journal, 26(3):557–573, 1979.

[3] George Cotsarelis, Tung-Tien Sun, and Robert M Lavker. Label-retaining cells reside in the bulge area of pilosebaceous unit: implications for follicular stem cells, hair cycle, and skin carcinogenesis. Cell, 61(7):1329–1337, 1990.

[4] Tudorita Tumbar, Geraldine Guasch, Valentina Greco, Cedric Blanpain, William E Lowry, Michael Rendl, and Elaine Fuchs. Defining the epithelial stem cell niche in skin. Science, 303(5656):359–363, 2004.

[5] Kai Kretzschmar and Fiona M Watt. Lineage tracing. Cell, 148(1):33–45, 2012.

[6] John E Sulston, Einhard Schierenberg, John G White, and J Nichol Thomson. The embryonic cell lineage of the nematode caenorhabditis elegans. Developmental biology, 100(1):64–119, 1983.

[7] Aden Forrow and Geoffrey Schiebinger. Lineageot is a unified framework for lineage tracing and trajectory inference. Nature communications, 12(1):4940, 2021.

[8] Geoffrey Schiebinger, Jian Shu, Marcin Tabaka, Brian Cleary, Vidya Subramanian, Aryeh Solomon, Joshua Gould, Siyan Liu, Stacie Lin, Peter Berube, et al. Optimal-transport analysis of single-cell gene expression identifies developmental trajectories in reprogramming. Cell, 176(4):928–943, 2019.

[9] Danilo Pellin, Mariana Loperfido, Cristina Baricordi, Samuel L Wolock, Annita Monte-peloso, Olga K Weinberg, Alessandra Biffi, Allon M Klein, and Luca Biasco. A comprehensive single cell transcriptional landscape of human hematopoietic progenitors. Nature communications, 10(1):2395, 2019.

[10] Betsabeh Khoramian Tusi, Samuel L Wolock, Caleb Weinreb, Yung Hwang, Daniel Hidalgo, Rapolas Zilionis, Ari Waisman, Jun R Huh, Allon M Klein, and Merav Socolovsky. Population snapshots predict early haematopoietic and erythroid hierarchies. Nature, 555(7694): 54–60, 2018.

[11] Jason D Buenrostro, M Ryan Corces, Caleb A Lareau, Beijing Wu, Alicia N Schep, Martin J Aryee, Ravindra Majeti, Howard Y Chang, and William J Greenleaf. Integrated single-cell analysis maps the continuous regulatory landscape of human hematopoietic differentiation. Cell, 173(6):1535–1548, 2018.

[12] Chengxiang Qiu, Beth K Martin, Ian C Welsh, Riza M Daza, Truc-Mai Le, Xingfan Huang, Eva K Nichols, Megan L Taylor, Olivia Fulton, Diana R O’Day, et al. A single-cell time-lapse of mouse prenatal development from gastrula to birth. Nature, pages 1–10, 2024.

[13] Sean C Bendall, Kara L Davis, El-ad David Amir, Michelle D Tadmor, Erin F Simonds, Tiffany J Chen, Daniel K Shenfeld, Garry P Nolan, and Dana Pe’er. Single-cell trajectory detection uncovers progression and regulatory coordination in human b cell development. Cell, 157(3):714–725, 2014.

[14] Cole Trapnell, Davide Cacchiarelli, Jonna Grimsby, Prapti Pokharel, Shuqiang Li, Michael Morse, Niall J Lennon, Kenneth J Livak, Tarjei S Mikkelsen, and John L Rinn. The dynamics and regulators of cell fate decisions are revealed by pseudotemporal ordering of single cells. Nature biotechnology, 32(4):381–386, 2014.

[15] Xiaojie Qiu, Qi Mao, Ying Tang, Li Wang, Raghav Chawla, Hannah A Pliner, and Cole Trapnell. Reversed graph embedding resolves complex single-cell trajectories. Nature methods, 14(10):979–982, 2017.

[16] F Alexander Wolf, Fiona K Hamey, Mireya Plass, Jordi Solana, Joakim S Dahlin, Berthold Göttgens, Nikolaus Rajewsky, Lukas Simon, and Fabian J Theis. Paga: graph abstraction reconciles clustering with trajectory inference through a topology preserving map of single cells. Genome biology, 20:1–9, 2019.

[17] Joshua D Welch, Alexander J Hartemink, and Jan F Prins. Slicer: inferring branched, non-linear cellular trajectories from single cell rna-seq data. Genome biology, 17:1–15, 2016.

[18] Kelly Street, Davide Risso, Russell B Fletcher, Diya Das, John Ngai, Nir Yosef, Elizabeth Purdom, and Sandrine Dudoit. Slingshot: cell lineage and pseudotime inference for single-cell transcriptomics. BMC genomics, 19:1–16, 2018.

[19] Huidong Chen, Luca Albergante, Jonathan Y Hsu, Caleb A Lareau, Giosuè Lo Bosco, Jihong Guan, Shuigeng Zhou, Alexander N Gorban, Daniel E Bauer, Martin J Aryee, et al. Single-cell trajectories reconstruction, exploration and mapping of omics data with stream. Nature communications, 10(1):1–14, 2019.

[20] Josip S Herman, null Sagar, and Dominic Gruen. Fateid infers cell fate bias in multipotent progenitors from single-cell rna-seq data. Nature methods, 15(5):379–386, 2018.

[21] Manu Setty, Michelle D Tadmor, Shlomit Reich-Zeliger, Omer Angel, Tomer Meir Salame, Pooja Kathail, Kristy Choi, Sean Bendall, Nir Friedman, and Dana Pe’er. Wishbone identifies bifurcating developmental trajectories from single-cell data. Nature biotechnology, 34(6): 637–645, 2016.

[22] Wouter Saelens, Robrecht Cannoodt, Helena Todorov, and Yvan Saeys. A comparison of single-cell trajectory inference methods. Nature biotechnology, 37(5):547–554, 2019.

[23] Daniel E Wagner and Allon M Klein. Lineage tracing meets single-cell omics: opportunities and challenges. Nature Reviews Genetics, 21(7):410–427, 2020.

[24] Lingfei Wang, Qian Zhang, Qian Qin, Nikolaos Trasanidis, Michael Vinyard, Huidong Chen, and Luca Pinello. Current progress and potential opportunities to infer single-cell developmental trajectory and cell fate. Current opinion in systems biology, 26:1–11, 2021.

[25] Louise Deconinck, Robrecht Cannoodt, Wouter Saelens, Bart Deplancke, and Yvan Saeys. Recent advances in trajectory inference from single-cell omics data. Current Opinion in Systems Biology, 27:100344, 2021.

[26] Daniel E Wagner, Caleb Weinreb, Zach M Collins, James A Briggs, Sean G Megason, and Allon M Klein. Single-cell mapping of gene expression landscapes and lineage in the zebrafish embryo. Science, 360(6392):981–987, 2018.

[27] Macy W Veling, Ye Li, Mike T Veling, Christopher Litts, Nigel Michki, Hao Liu, Bing Ye, and Dawen Cai. Identification of neuronal lineages in the drosophila peripheral nervous system with a “digital” multi-spectral lineage tracing system. Cell reports, 29(10):3303–3312, 2019.

[28] Michelle M Chan, Zachary D Smith, Stefanie Grosswendt, Helene Kretzmer, Thomas M Norman, Britt Adamson, Marco Jost, Jeffrey J Quinn, Dian Yang, Matthew G Jones, et al. Molecular recording of mammalian embryogenesis. Nature, 570(7759):77–82, 2019.

[29] Adriano Bolondi, Benjamin K Law, Helene Kretzmer, Seher Ipek Gassaloglu, René Buschow, Christina Riemenschneider, Dian Yang, Maria Walther, Jesse V Veenvliet, Alexander Meissner, et al. Reconstructing axial progenitor field dynamics in mouse stem cell-derived embryoids. Developmental cell, 2024.

[30] Caleb Weinreb, Alejo Rodriguez-Fraticelli, Fernando D Camargo, and Allon M Klein. Lineage tracing on transcriptional landscapes links state to fate during differentiation. Science, 367(6479):eaaw3381, 2020.

[31] Zhisong He, Ashley Maynard, Akanksha Jain, Tobias Gerber, Rebecca Petri, Hsiu-Chuan Lin, Malgorzata Santel, Kevin Ly, Jean-Samuel Dupré, Leila Sidow, et al. Lineage recording in human cerebral organoids. Nature methods, 19(1):90–99, 2022.

[32] Wenjun Kong, Brent A Biddy, Kenji Kamimoto, Junedh M Amrute, Emily G Butka, and Samantha A Morris. Celltagging: combinatorial indexing to simultaneously map lineage and identity at single-cell resolution. Nature protocols, 15(3):750–772, 2020.

[33] Kunal Jindal, Mohd Tayyab Adil, Naoto Yamaguchi, Xue Yang, Helen C Wang, Kenji Kamimoto, Guillermo C Rivera-Gonzalez, and Samantha A Morris. Single-cell lineage capture across genomic modalities with celltag-multi reveals fate-specific gene regulatory changes. Nature Biotechnology, pages 1–14, 2023.

[34] Aaron McKenna, Gregory M Findlay, James A Gagnon, Marshall S Horwitz, Alexander F Schier, and Jay Shendure. Whole-organism lineage tracing by combinatorial and cumulative genome editing. Science, 353(6298):aaf7907, 2016.

[35] Bushra Raj, James A Gagnon, and Alexander F Schier. Large-scale reconstruction of cell lineages using single-cell readout of transcriptomes and crispr–cas9 barcodes by scgestalt. Nature protocols, 13(11):2685–2713, 2018.

[36] Anna Alemany, Maria Florescu, Chloé S Baron, Josi Peterson-Maduro, and Alexander Van Oudenaarden. Whole-organism clone tracing using single-cell sequencing. Nature, 556 (7699):108–112, 2018.

[37] Bastiaan Spanjaard, Bo Hu, Nina Mitic, Pedro Olivares-Chauvet, Sharan Janjuha, Nikolay Ninov, and Jan Philipp Junker. Simultaneous lineage tracing and cell-type identification using crispr–cas9-induced genetic scars. Nature biotechnology, 36(5):469–473, 2018.

[38] Jeffrey J Quinn, Matthew G Jones, Ross A Okimoto, Shigeki Nanjo, Michelle M Chan, Nir Yosef, Trever G Bivona, and Jonathan S Weissman. Single-cell lineages reveal the rates, routes, and drivers of metastasis in cancer xenografts. Science, 371(6532):eabc1944, 2021.

[39] Dian Yang, Matthew G Jones, Santiago Naranjo, William M Rideout, Kyung Hoi Joseph Min, Raymond Ho, Wei Wu, Joseph M Replogle, Jennifer L Page, Jeffrey J Quinn, et al. Lineage tracing reveals the phylodynamics, plasticity, and paths of tumor evolution. Cell, 185(11): 1905–1923, 2022.

[40] Matthew G Jones, Alex Khodaverdian, Jeffrey J Quinn, Michelle M Chan, Jeffrey A Hussmann, Robert Wang, Chenling Xu, Jonathan S Weissman, and Nir Yosef. Inference of single-cell phylogenies from lineage tracing data using cassiopeia. Genome biology, 21(1):1–27, 2020.

[41] Palash Sashittal, Henri Schmidt, Michelle M Chan, and Benjamin J Raphael. Startle: a star homoplasy approach for crispr-cas9 lineage tracing. bioRxiv, pages 2022–12, 2022.

[42] Xinhai Pan, Hechen Li, Pranav Putta, and Xiuwei Zhang. Linrace: cell division history reconstruction of single cells using paired lineage barcode and gene expression data. Nature Communications, 14(1):8388, 2023.

[43] Jean Feng, William S Dewitt III, Aaron McKenna, Noah Simon, Amy D Willis, and FREDERICK A MATSEN IV. Estimation of cell lineage trees by maximum-likelihood phylogenetics. The annals of applied statistics, 15(1):343, 2021.

[44] Wuming Gong, Alejandro A Granados, Jingyuan Hu, Matthew G Jones, Ofir Raz, Irepan Salvador-Martínez, Hanrui Zhang, Ke-Huan K Chow, Il-Youp Kwak, Renata Retkute, et al. Benchmarked approaches for reconstruction of in vitro cell lineages and in silico models of c. elegans and m. musculus developmental trees. Cell systems, 12(8):810–826, 2021.

[45] Hamim Zafar, Chieh Lin, and Ziv Bar-Joseph. Single-cell lineage tracing by integrating crispr-cas9 mutations with transcriptomic data. Nature communications, 11(1):3055, 2020.

[46] Uyen Mai, Gillian Chu, and Benjamin J Raphael. Maximum likelihood inference of time-scaled cell lineage trees with mixed-type missing data. In International Conference on Research in Computational Molecular Biology, pages 360–363. Springer, 2024.

[47] Weixiang Fang, Claire M Bell, Abel Sapirstein, Soichiro Asami, Kathleen Leeper, Donald J Zack, Hongkai Ji, and Reza Kalhor. Quantitative fate mapping: A general framework for analyzing progenitor state dynamics via retrospective lineage barcoding. Cell, 185(24):4604–4620, 2022.

[48] Jesse V Veenvliet, Adriano Bolondi, Helene Kretzmer, Leah Haut, Manuela Scholze-Wittler, Dennis Schifferl, Frederic Koch, Léo Guignard, Abhishek Sampath Kumar, Milena Pustet, et al. Mouse embryonic stem cells self-organize into trunk-like structures with neural tube and somites. Science, 370(6522):eaba4937, 2020.

[49] Adriano Bolondi, Leah Haut, Seher Ipek Gassaloglu, Polly Burton, Helene Kretzmer, René Buschow, Alexander Meissner, Bernhard G Herrmann, and Jesse V Veenvliet. Generation of mouse pluripotent stem cell-derived trunk-like structures: an in vitro model of post-implantation embryogenesis. Bio-protocol, 11(11):e4042–e4042, 2021.

[50] Willy Feller. Die grundlagen der volterraschen theorie des kampfes ums dasein in wahrscheinlichkeitstheoretischer behandlung. Acta Biotheoretica, 5(1):11–40, 1939.

[51] Paul Jaccard. The distribution of the flora in the alpine zone. 1. New phytologist, 11(2): 37–50, 1912.

[52] Kun Wang, Liangzhen Hou, Xin Wang, Xiangwei Zhai, Zhaolian Lu, Zhike Zi, Weiwei Zhai, Xionglei He, Christina Curtis, D. Zhou, et al. Phylovelo enhances transcriptomic velocity field mapping using monotonically expressed genes. Nature Biotechnology, pages 1–12, 2023.

[53] Kirstie A Lawson, Juanito J Meneses, and Roger A Pedersen. Clonal analysis of epiblast fate during germ layer formation in the mouse embryo. Development, 113(3):891–911, 1991.

[54] Sylvie Forlani, Kirstie A Lawson, and Jacqueline Deschamps. Acquisition of hox codes during gastrulation and axial elongation in the mouse embryo. 2003.

[55] Tatiana Solovieva, Valerie Wilson, and Claudio D Stern. A niche for axial stem cells-a cellular perspective in amniotes. Developmental Biology, 490:13–21, 2022.

[56] Mounia Lagha, Silvia Brunelli, Graziella Messina, Ana Cumano, Tsutomu Kume, Frédéric Relaix, and Margaret E Buckingham. Pax3: Foxc2 reciprocal repression in the somite modulates muscular versus vascular cell fate choice in multipotent progenitors. Developmental cell, 17(6):892–899, 2009.

[57] Phong Dang Nguyen, Georgina Elizabeth Hollway, Carmen Sonntag, Lee Barry Miles, Thomas Edward Hall, Silke Berger, Kristine Joy Fernandez, David Baruch Gurevich, Nicholas James Cole, Sara Alaei, et al. Haematopoietic stem cell induction by somite-derived endothelial cells controlled by meox1. Nature, 512(7514):314–318, 2014.

[58] Caleb Weinreb, Samuel Wolock, and Allon M Klein. Spring: a kinetic interface for visualizing high dimensional single-cell expression data. Bioinformatics, 34(7):1246–1248, 2018.

[59] Jun Seita and Irving L Weissman. Hematopoietic stem cell: self-renewal versus differentiation. Wiley Interdisciplinary Reviews: Systems Biology and Medicine, 2(6):640–653, 2010.

[60] Koichi Akashi, David Traver, Toshihiro Miyamoto, and Irving L Weissman. A clonogenic common myeloid progenitor that gives rise to all myeloid lineages. Nature, 404(6774):193–197, 2000.

[61] Fang Dong, Sha Hao, Sen Zhang, Caiying Zhu, Hui Cheng, Zining Yang, Fiona K Hamey, Xiaofang Wang, AI Gao, Fengjiao Wang, et al. Differentiation of transplanted haematopoietic stem cells tracked by single-cell transcriptomic analysis. Nature cell biology, 22(6):630–639, 2020.

[62] Donald Metcalf. Hematopoietic cytokines. Blood, The Journal of the American Society of Hematology, 111(2):485–491, 2008.

[63] Balyn W Zaro, Joseph J Noh, Victoria L Mascetti, Janos Demeter, Benson George, Monika Zukowska, Gunsagar S Gulati, Rahul Sinha, Ryan A Flynn, Allison Banuelos, et al. Proteomic analysis of young and old mouse hematopoietic stem cells and their progenitors reveals post-transcriptional regulation in stem cells. Elife, 9:e62210, 2020.

[64] Hui Cheng, Zhaofeng Zheng, and Tao Cheng. New paradigms on hematopoietic stem cell differentiation. Protein & cell, 11(1):34–44, 2020.

[65] Joana Carrelha, Yiran Meng, Laura M Kettyle, Tiago C Luis, Ruggiero Norfo, Verónica Alcolea, Hanane Boukarabila, Francesca Grasso, Adriana Gambardella, Amit Grover, et al. Hierarchically related lineage-restricted fates of multipotent haematopoietic stem cells. Nature, 554(7690):106–111, 2018.

[66] Ryo Yamamoto, Yohei Morita, Jun Ooehara, Sanae Hamanaka, Masafumi Onodera, Karl Lenhard Rudolph, Hideo Ema, and Hiromitsu Nakauchi. Clonal analysis unveils self-renewing lineage-restricted progenitors generated directly from hematopoietic stem cells. Cell, 154(5):1112–1126, 2013.

[67] David F Robinson and Leslie R Foulds. Comparison of phylogenetic trees. Mathematical biosciences, 53(1-2):131–147, 1981.

[68] Jinyang Li and Ben Z Stanger. How tumor cell dedifferentiation drives immune evasion and resistance to immunotherapy. Cancer research, 80(19):4037–4041, 2020.

[69] Stewart Sell. Cellular origin of cancer: dedifferentiation or stem cell maturation arrest? Environmental health perspectives, 101(suppl 5):15–26, 1993.

[70] Yosuke Yamada, Hironori Haga, and Yasuhiro Yamada. Concise review: dedifferentiation meets cancer development: proof of concept for epigenetic cancer. Stem cells translational medicine, 3(10):1182–1187, 2014.

[71] Dinorah Friedmann-Morvinski and Inder M Verma. Dedifferentiation and reprogramming: origins of cancer stem cells. EMBO reports, 15(3):244–253, 2014.

[72] Ke-Huan K Chow, Mark W Budde, Alejandro A Granados, Maria Cabrera, Shinae Yoon, Soomin Cho, Ting-Hao Huang, Noushin Koulena, Kirsten L Frieda, Long Cai, et al. Imaging cell lineage with a synthetic digital recording system. Science, 372(6538):eabb3099, 2021.

[73] Duncan M Chadly, Kirsten L Frieda, Chen Gui, Leslie Klock, Martin Tran, Margaret Y Sui, Yodai Takei, Remco Bouckaert, Carlos Lois, Long Cai, et al. Reconstructing cell histories in space with image-readable base editor recording. bioRxiv, pages 2024–01, 2024.

[74] Li Li, Sarah Bowling, Sean E McGeary, Qi Yu, Bianca Lemke, Karel Alcedo, Yuemeng Jia, Xugeng Liu, Mark Ferreira, Allon M Klein, et al. A mouse model with high clonal barcode diversity for joint lineage, transcriptomic, and epigenomic profiling in single cells. Cell, 186 (23):5183–5199, 2023.

[75] Walter M Fitch. Toward defining the course of evolution: minimum change for a specific tree topology. Systematic Biology, 20(4):406–416, 1971.

[76] David Sankoff and Pascale Rousseau. Locating the vertices of a steiner tree in an arbitrary metric space. Mathematical Programming, 9(1):240–246, 1975.

[77] Gurobi Optimization, LLC. Gurobi Optimizer Reference Manual, 2024. URL https://www.gurobi.com.

[78] Ville Satopaa, Jeannie Albrecht, David Irwin, and Barath Raghavan. Finding a” kneedle” in a haystack: Detecting knee points in system behavior. In 2011 31st international conference on distributed computing systems workshops, pages 166–171. IEEE, 2011.

[79] Derek JS Robinson. An introduction to abstract algebra. Walter de Gruyter, 2003.

[80] Robert R Sokal and Charles D Michener. A statistical method for evaluating systematic relationships. 1958.

[81] Xiaojie Qiu, Yan Zhang, Jorge D Martin-Rufino, Chen Weng, Shayan Hosseinzadeh, Dian Yang, Angela N Pogson, Marco Y Hein, Kyung Hoi Joseph Min, Li Wang, et al. Mapping transcriptomic vector fields of single cells. Cell, 185(4):690–711, 2022.

[82] Markus Chimani, Sven Rahmann, and Sebastian Böcker. Exact ilp solutions for phylogenetic minimum flip problems. In Proceedings of the First ACM International Conference on Bioinformatics and Computational Biology, pages 147–153, 2010.

[83] Dan Gusfield. Recombinatorics: the algorithmics of ancestral recombination graphs and explicit phylogenetic networks.

[84] Dan Gusfield. Efficient algorithms for inferring evolutionary trees. Networks, 21(1):19–28, 1991.

[85] Duhong Chen, Oliver Eulenstein, David Fernández-Baca, and Michael Sanderson. Supertrees by flipping. In International Computing and Combinatorics Conference, pages 391–400. Springer, 2002.

[86] Motoo Kimura. The number of heterozygous nucleotide sites maintained in a finite population due to steady flux of mutations. Genetics, 61(4):893, 1969.

[87] Erfan Sadeqi Azer, Farid Rashidi Mehrabadi, Salem Malikić, Xuan Cindy Li, Osnat Bartok, Kevin Litchfield, Ronen Levy, Yardena Samuels, Alejandro A Schäffer, E Michael Gertz, et al. Phiscs-bnb: a fast branch and bound algorithm for the perfect tumor phylogeny reconstruction problem. Bioinformatics, 36(Supplement 1):i169–i176, 2020.

[88] Duhong Chen, Oliver Eulenstein, David Fernandez-Baca, and Michael Sanderson. Minimum-flip supertrees: complexity and algorithms. IEEE/ACM transactions on computational biology and bioinformatics, 3(2):165–173, 2006.

[89] Morris Goodman, John Czelusniak, G William Moore, Alejo E Romero-Herrera, and Genji Matsuda. Fitting the gene lineage into its species lineage, a parsimony strategy illustrated by cladograms constructed from globin sequences. Systematic Biology, 28(2):132–163, 1979.

[90] Richard M Karp. Reducibility among combinatorial problems. Springer, 2010.

[91] Catherine Greenhill. The complexity of counting colourings and independent sets in sparse graphs and hypergraphs. computational complexity, 9:52–72, 2000.

